## Supplementary material for "Striking parallels between dorsoventral patterning in *Drosophila* and *Gryllus* reveal a complex evolutionary history behind a model gene regulatory network": Pechmann et al Supplement

#### 1. Videos and supplementary videos

**Video 1.** Mesoderm internalization.
Time lapse imaging using the pXLBGact Histone2B:eGFP transgenic *Gryllus* line (Nakamura et al., 2010). Ventral surface views and z-sections from early to late gem anlage condensation (ES 2.2- ES2.6). Staging according to (Donoughe and Extavour, 2016; Sarashina et al., 2005).

**Video 2.** Early development of an embryo lacking BMP signalling (*Gb-dpp2* KD).
Time lapse imaging using the pXLBGact Histone2B:eGFP transgenic *Gryllus* line (Nakamura et al., 2010). The movie shows the development of a control and a *Gb-dpp2* RNAi embryo from uniform blastoderm stage (egg stage 2; embryonic stage 1.5) until embryonic stage 5 (staging according to (Donoughe and Extavour, 2016)). The germ-rudiment condenses ventrally in the control embryo (15h-30h), the serosa closes over the embryo (35h-40h) and anatrepsis takes place (40h onwards). The germ-rudiment condenses towards the ventral-posterior in the *Gb-dpp2* knockdown embryos and ectopic tissue folding is taking place (40h onwards).

**Video 3.** Complete development of embryo lacking BMP signalling (*Gb-tkv* KD).
Six days of constant live imaging using the pXLBGact Histone2B:eGFP transgenic *Gryllus* line (Nakamura et al., 2010). The movie shows the development of a *Gb-tkv* RNAi embryo from uniform blastoderm stage (egg stage 2, staging according to (Donoughe and Extavour, 2016)) until embryonic stage 11. The malformed *Gb-tkv* RNAi embryo is released from the yolk and the serosa at the end of the movie (132h onwards).

**Video 4.** Development of embryos lacking Toll signalling (*Gb-Toll1* KD).
Time lapse imaging using the pXLBGact Histone2B:eGFP transgenic *Gryllus* line (Nakamura et al., 2010). The movie shows the development of a control and three *Gb-Toll1* RNAi embryos from uniform blastoderm stage (egg stage 2; embryonic stage 1.5) until embryonic stage 6-7 (staging according to (Donoughe and Extavour, 2016)). The germ-rudiment condenses ventrally in the control embryo (10h-30h), the serosa closes over the embryo (30h-40h) and anatrepsis takes place (35h onwards). The germ-rudiment condenses

towards the posterior in the *Gb-Toll1* knockdown embryos (20h-35h). However, the posterior cap of cells sinks into the yolk during anatrepsis (35h onwards).

**Video S1.** Early development of embryo lacking BMP signalling (*Gb-dpp2* KD).
Time lapse imaging using the pXLBGact Histone2B:eGFP transgenic *Gryllus* line (Nakamura et al., 2010). The movie shows the development of a *Gb-dpp2* RNAi embryo from early germ anlage condensation (ES 2,2) until ES 5. The border between serosa and germ anlage lacks DV asymmetry. The germ anlage condenses symmetrically towards the posterior pole. Staging according to (Donoughe and Extavour, 2016; Sarashina et al., 2005)).

**Video S2.** Complete development of embryos lacking Toll signalling (*Gb-Toll1* KD).
Time lapse imaging of a control and a *Gb-Toll1* RNAi embryo from egg stage 8 onwards. The serosa detaches from the posterior pole in both embryos (starting at 20 h). While the control embryo undergoes katatrepsis (starting at 53 h) the serosa of the *Gb-Toll1* RNAi embryo also detaches from the anterior (starting at 72 h). This leads to a compaction of the yolk within the centre of the egg. This represents the strong phenotype that is observed after the knockdown of *Gb-Toll1*, *Gb-dl1 and Gb-spz*. Yolk is taken up into the gut in the control embryo (starting at 110 h) and the movie ends at egg stage 19. Staging according to (Donoughe and Extavour, 2016).

#### 2. Supplementary Figures

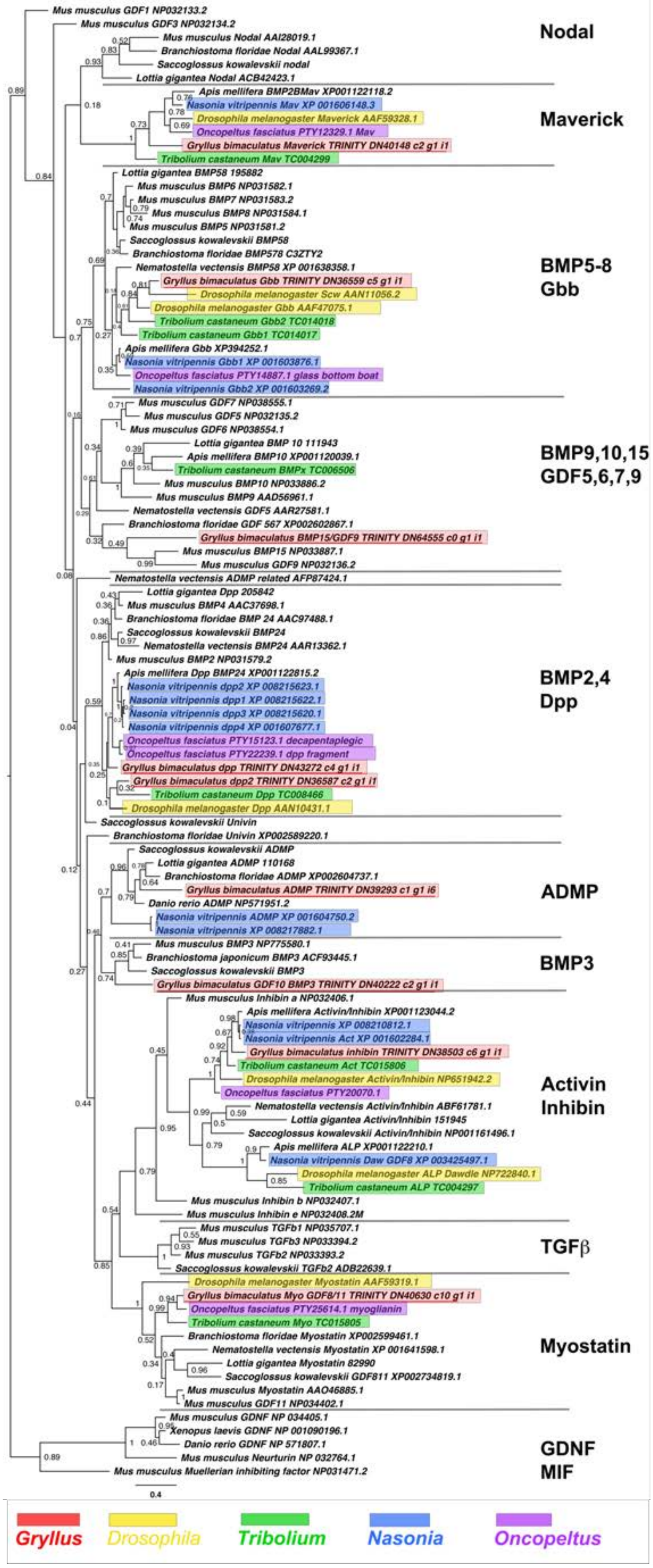

**Figure S1.** Phylogeny of BMP and TGFβ-like ligand class interrelationships across the Metazoa, as determined by Bayesian (Huelsenbeck and Ronquist 2001) methods. Alignment generated using MAFFT –add (Katoh and Standley 2013) with the previously established dataset used in Kenny et al 2015 and homologues of known identity from NCBI's nr database using the L-INS-i strategy resulting in a 61 informative amino acid alignment of the mature ligand domains after the exclusion of gaps. Phylogeny determined using the WAG model (Whelan and Goldman 2001). Posterior probabilities (after 5,000,000 generations) can be seen at the nodes of trees. Phylogeny rooted with known Neurturin and GDNF outgroups. Scale bars represent substitutions per site at given distances. The sequences of *Gryllus bimaculatus*, *Drosophila melanogaster*, *Tribolium castaneum*, *Nasonia vitripennis* and *Oncopeltus fasciatus* are highlighted by indicated colours.

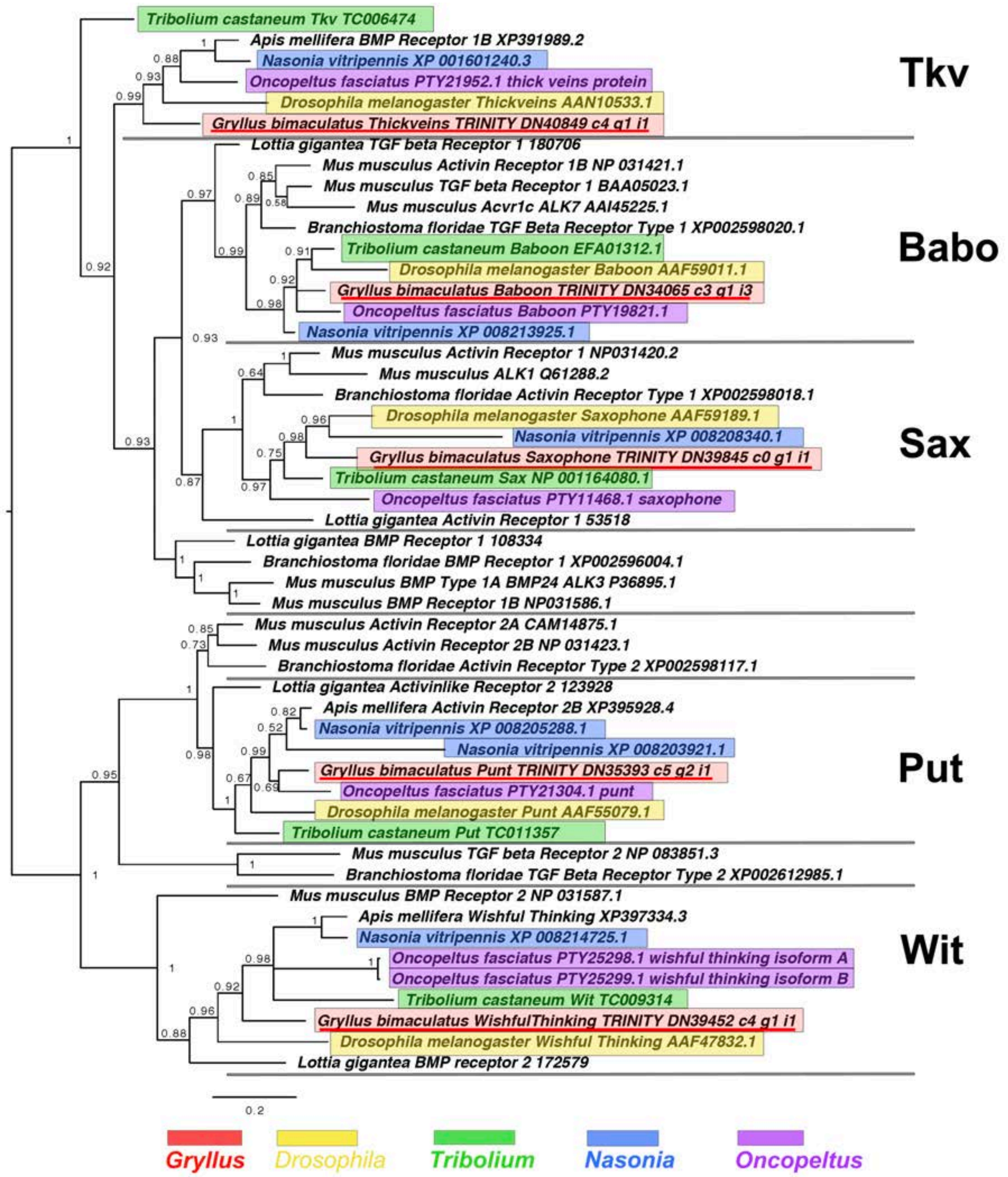


**Figure S2.** TGFβ and BMP receptor molecule interrelationships across the Metazoa, as determined by Bayesian (Huelsenbeck and Ronquist 2001) methods, and rooted at the midpoint.
Predominantly TGFβ-like and BMP-like cascade receptors shown in green and blue respectively. Alignment generated using MAFFT –add (Katoh and Standley 2013) and homologues of known identity from NCBI's nr database against the previously established dataset used in Kenny et al 2015 using the G-iNS-i strategy resulting in a 136 informative amino acid alignment spanning the protein kinase domain (PFAM PF00069). Phylogenies determined using the WAG model (Whelan and Goldman 2001). Posterior probabilities can be seen at nodes. Scale bars represent substitutions per site at given distances. The sequences of *Gryllus bimaculatus*, *Drosophila melanogaster*, *Tribolium castaneum*, *Nasonia vitripennis* and *Oncopeltus fasciatus* are highlighted by indicated colours.

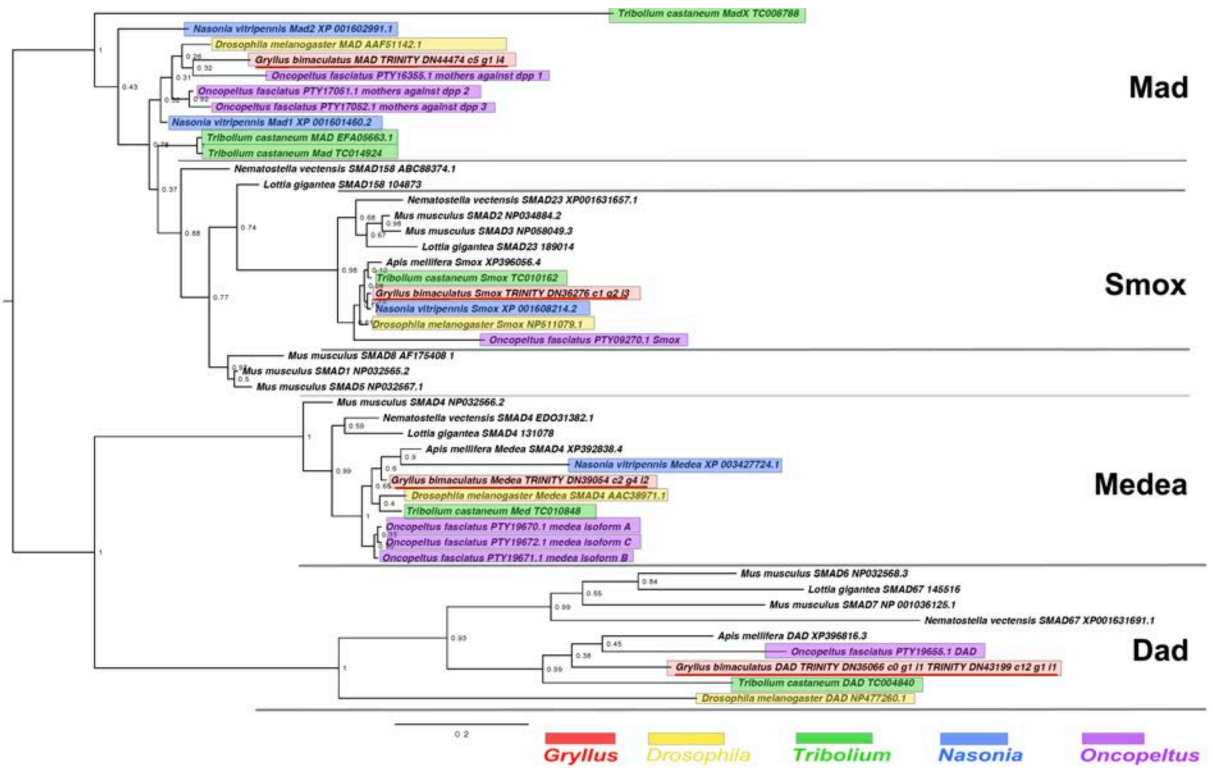


**Figure S3.** Smad and Dad interrelationships across the Metazoa, as determined by Bayesian (Huelsenbeck and Ronquist 2001) methods.
Alignment generated using MAFFT –add (Katoh and Standley 2013) against the previously established dataset used in Kenny et al 2015 and homologues of known identity from NCBI's nr database using the G-iNS-i strategy, with the section used for analysis a 127 informative amino acid region spanning the MH2 domain (Pfam PF03166). Phylogenies determined using the Jones model, rooted at midpoint. Posterior probabilities (after 2 million generations) can be seen at nodes. Scale bars represent substitutions per site at given distances. The sequences of *Gryllus bimaculatus*, *Drosophila melanogaster*, *Tribolium castaneum*, *Nasonia vitripennis* and *Oncopeltus fasciatus* are highlighted by indicated colours.

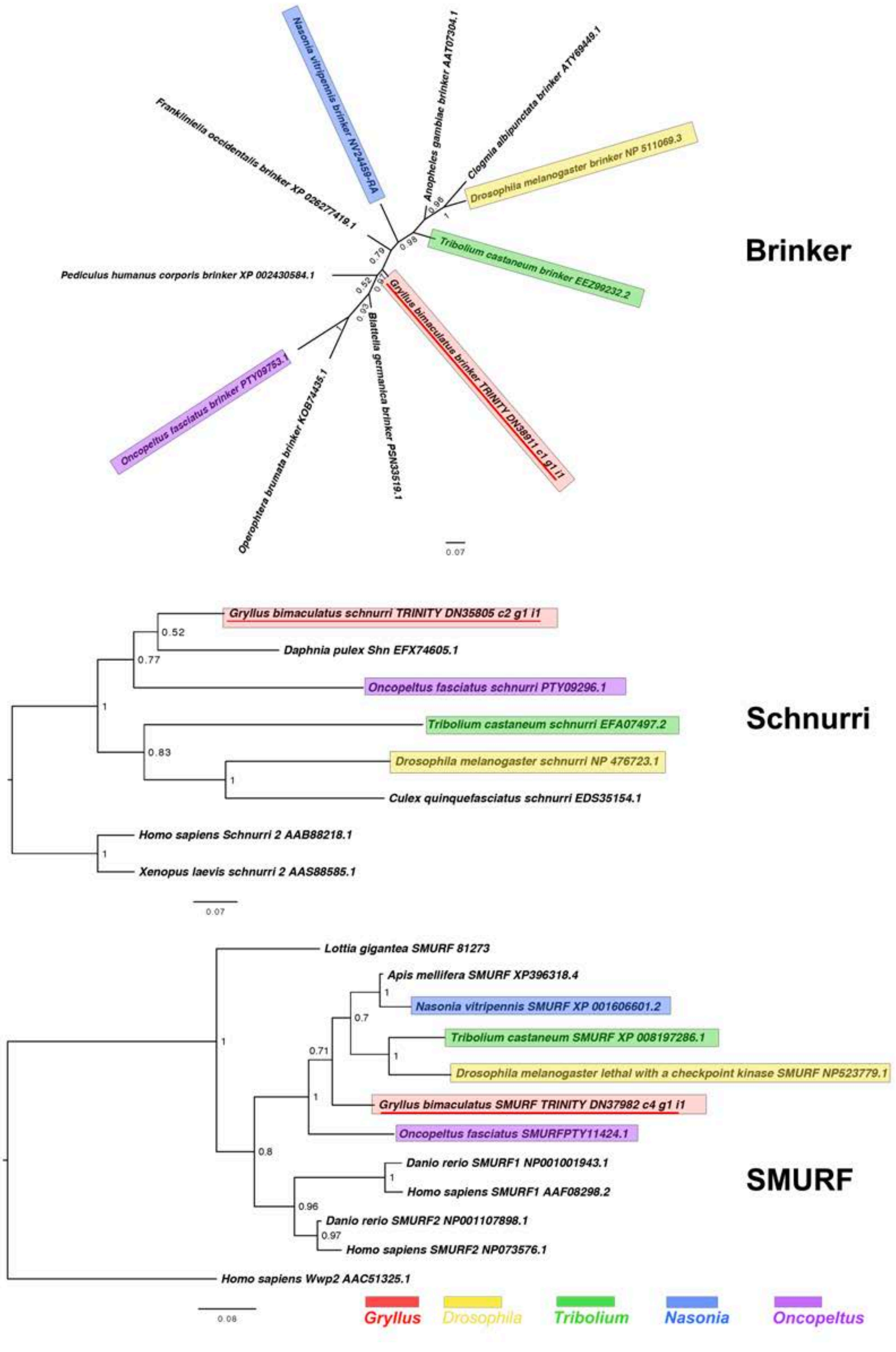


**Figure S4.** Brinker, Schnurri and SMURF interrelationships as determined by Bayesian (Huelsenbeck and Ronquist 2001) methods.
Brinker phylogenies inferred on the basis of a MAFFT alignment (Katoh and Standley 2013, G-INS-i strategy vs homologues of known identity from NCBI's nr database) with 66 informative sites, analysed under the Dayhoff model, and shown unrooted. Schnurri phylogeny based on a 135 informative amino acid global alignment generated using MAFFT (Katoh and Standley 2013) alongside homologues of known identity from NCBI's nr database using the G-iNS-i strategy, analysed under the WAG model, and rooted with known vertebrate sequence. The Smurf phylogeny was calculated under the Jones model, based on a

195 informative amino acid alignment generated using MAFFT – add (Katoh and Standley 2013) using the G-iNS-i strategy, against the Kenny et al dataset and homologues of known identity from NCBI's nr database with outgroup specified as *H. sapiens* Wwp2 (AAC51325.1). Scale bars represent substitutions per site at given distances. The sequences of *Gryllus bimaculatus*, *Drosophila melanogaster*, *Tribolium castaneum*, *Nasonia vitripennis* and *Oncopeltus fasciatus* are highlighted by indicated colours.

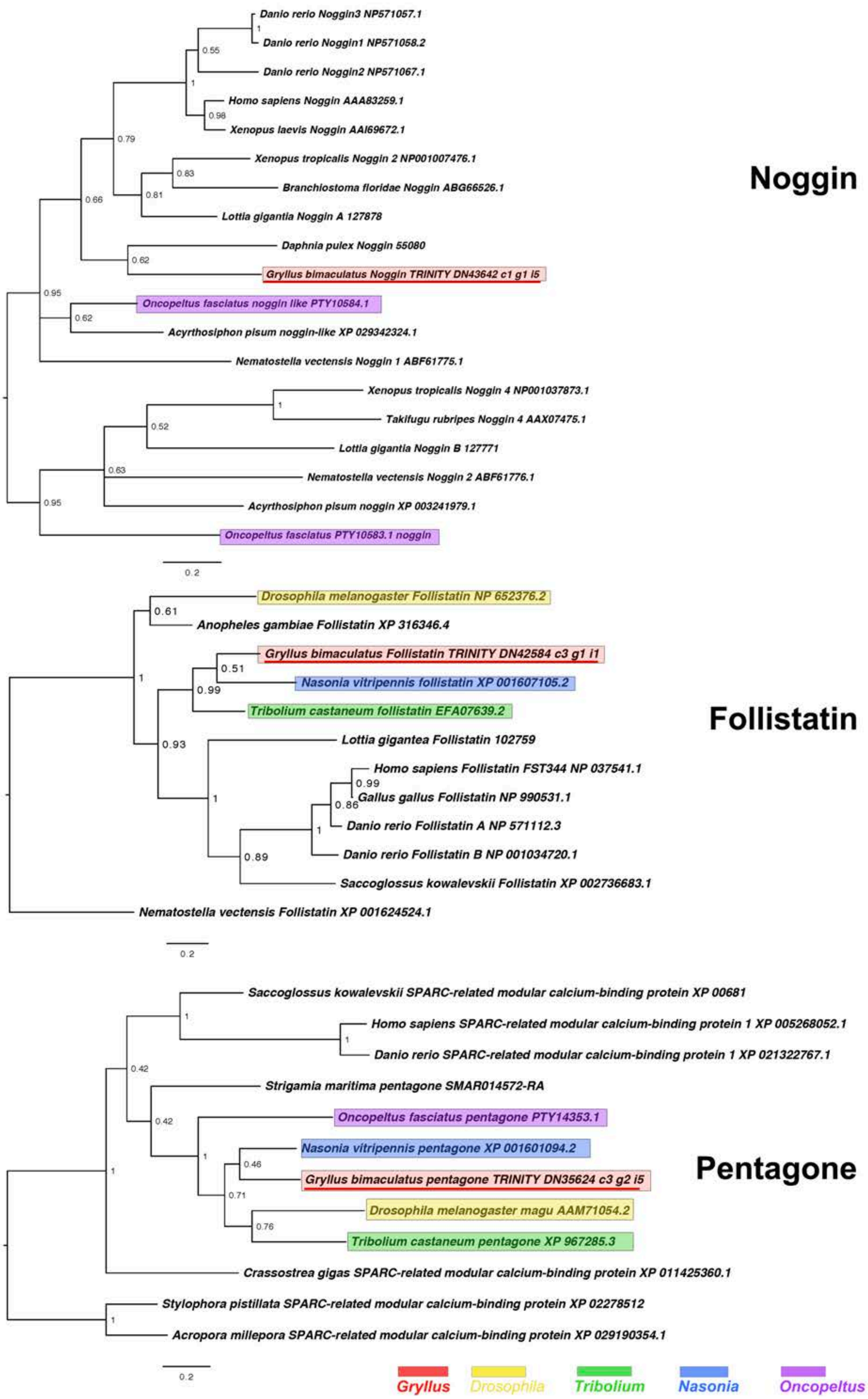
Danio rerio Noggin3 NP571057.1
Danio rerio Noggin1 NP571058.2
Danio rerio Noggin2 NP571067.1
Homo sapiens Noggin AAA83259.1
Xenopus laevis Noggin AAI69672.1
Xenopus tropicalis Noggin 2 NP001007476.1
Branchiostoma floridae Noggin ABG66526.1
Lottia gigantia Noggin A 127878
Daphnia pulex Noggin 55080
Gryllus bimaculatus Noggin TRINITY DN43642 c1 g1 i5
Oncopeltus fasciatus noggin like PTY10584.1
Acyrthosiphon pisum noggin-like XP 029342324.1
Nematostella vectensis Noggin 1 ABF61775.1
Xenopus tropicalis Noggin 4 NP001037873.1
Takifugu rubripes Noggin 4 AAX07475.1
Lottia gigantia Noggin B 127771
Nematostella vectensis Noggin 2 ABF61776.1
Acyrthosiphon pisum noggin XP 003241979.1
Oncopeltus fasciatus PTY10583.1 noggin
Noggin
0.2
Drosophila melanogaster Follistatin NP 652376.2
Anopheles gambiae Follistatin XP 316346.4
Gryllus bimaculatus Follistatin TRINITY DN42584 c3 g1 i1
Nasonia vitripennis follistatin XP 001607105.2
Tribolium castaneum follistatin EFA07639.2
Lottia gigantea Follistatin 102759
Homo sapiens Follistatin FST344 NP 037541.1
Gallus gallus Follistatin NP 990531.1
Danio rerio Follistatin A NP 571112.3
Danio rerio Follistatin B NP 001034720.1
Saccoglossus kowalevskii Follistatin XP 002736683.1
Nematostella vectensis Follistatin XP 001624524.1
Follistatin
0.2
Saccoglossus kowalevskii SPARC-related modular calcium-binding protein XP 00681
Homo sapiens SPARC-related modular calcium-binding protein 1 XP 005268052.1
Danio rerio SPARC-related modular calcium-binding protein 1 XP 021322767.1
Strigamia maritima pentagone SMAR014572-RA
Oncopeltus fasciatus pentagone PTY14353.1
Nasonia vitripennis pentagone XP 001601094.2
Gryllus bimaculatus pentagone TRINITY DN35624 c3 g2 i5
Drosophila melanogaster magu AAM71054.2
Tribolium castaneum pentagone XP 967285.3
Crassostrea gigas SPARC-related modular calcium-binding protein XP 011425360.1
Stylophora pistillata SPARC-related modular calcium-binding protein XP 02278512
Acropora millepora SPARC-related modular calcium-binding protein XP 029190354.1
Pentagone
0.2
Gryllus
Drosophila
Tribolium
Nasonia
Oncopeltus

**Figure S5**. Noggin, Follistatin and Pentagone interrelationships across the Metazoa, as determined using Bayesian (Huelsenbeck and Ronquist 2001) methods.
Noggin phylogeny based on a 80 informative amino acid global alignment generated using MAFFT -add (Katoh and Standley 2013) with the previously established dataset used in Kenny et al 2015 and homologues of known identity from NCBI's nr database using the G-iNS-i strategy rooted at midpoint. Posterior probabilities (2 million generations) can be seen at the base of nodes. Follistatin phylogenies inferred on the basis of a MAFFT -add alignment (Katoh and Standley 2013, G-INS-i strategy vs the Kenny et al 2015 dataset and homologues of known identity from NCBI's nr database) with 149 informative sites, analysed under the WAG model (Whelan and Goldman 2001), rooted with *N. vectensis* Follistatin (XP 001624524.1). Posterior probabilities can be seen at the base of nodes. Pentagone phylogeny based on a 220 informative amino acid global alignment generated using MAFFT (Katoh and Standley 2013) alongside homologues of known identity from NCBI's nr database using the G-iNS-i strategy, rooted with cnidarian SPARC-related modular calcium-binding protein sequence. Tree was run for 2 million generations under the WAG model. Posterior probabilities can be seen at the base of nodes. Scale bars represent substitutions per site at given distances. The sequences of *Gryllus bimaculatus*, *Drosophila melanogaster*, *Tribolium castaneum*, *Nasonia vitripennis* and *Oncopeltus fasciatus* are highlighted by indicated colours.

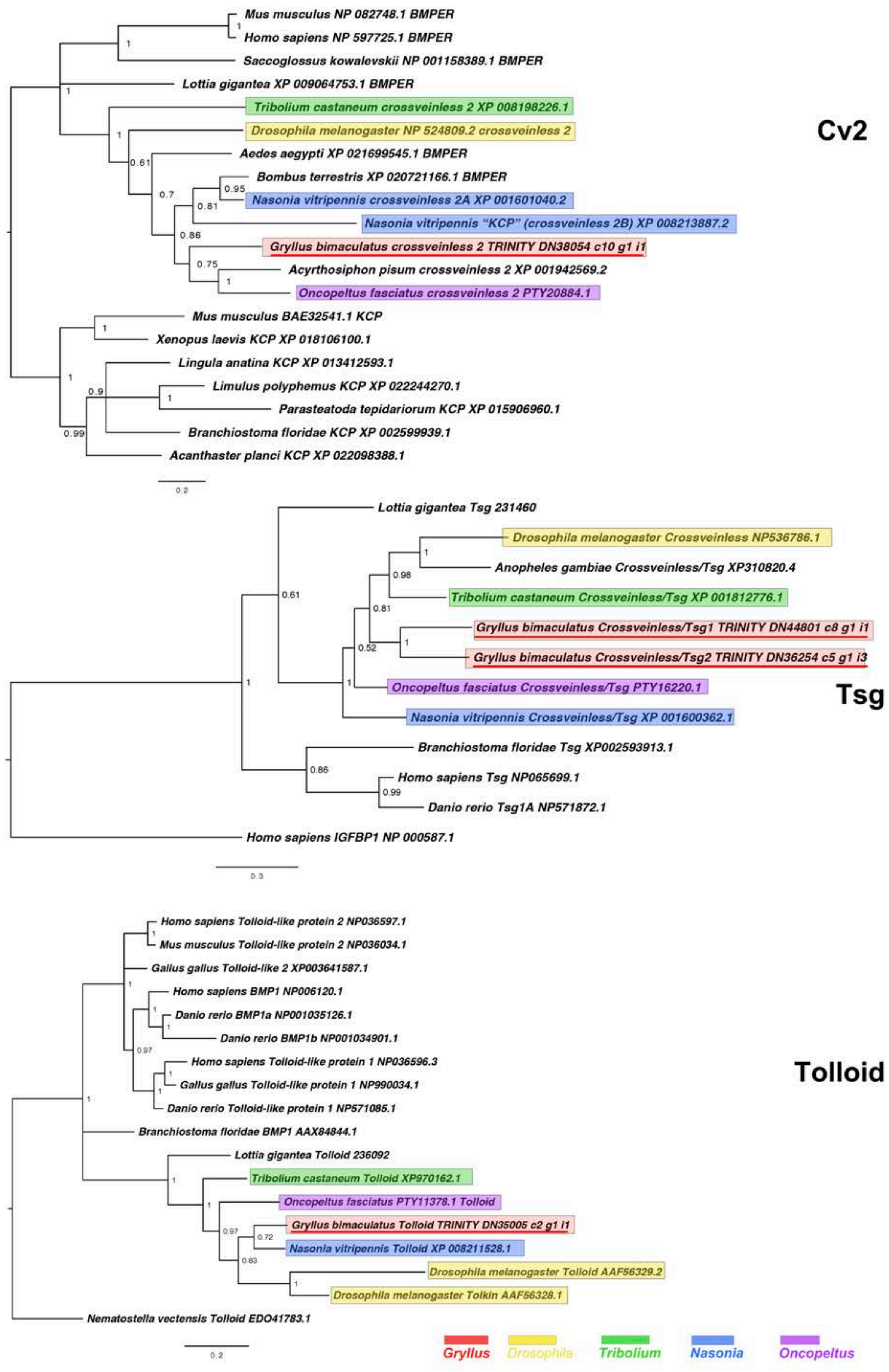
Cv2
Mus musculus NP 082748.1 BMPER
Homo sapiens NP 597725.1 BMPER
Saccoglossus kowalevskii NP 001158389.1 BMPER
Lottia gigantea XP 009064753.1 BMPER
Tribolium castaneum crossveinless 2 XP 008198226.1
Drosophila melanogaster NP 524809.2 crossveinless 2
Aedes aegypti XP 021699545.1 BMPER
Bombus terrestris XP 020721166.1 BMPER
Nasonia vitripennis crossveinless 2A XP 001601040.2
Nasonia vitripennis "KCP" (crossveinless 2B) XP 008213887.2
Gryllus bimaculatus crossveinless 2 TRINITY DN38054 c10 g1 i1
Acyrthosiphon pisum crossveinless 2 XP 001942569.2
Oncopeltus fasciatus crossveinless 2 PTY20884.1
Mus musculus BAE32541.1 KCP
Xenopus laevis KCP XP 018106100.1
Lingula anatina KCP XP 013412593.1
Limulus polyphemus KCP XP 022244270.1
Parasteatoda tepidariorum KCP XP 015906960.1
Branchiostoma floridae KCP XP 002599939.1
Acanthaster planci KCP XP 022098388.1
0.2
Tsg
Lottia gigantea Tsg 231460
Drosophila melanogaster Crossveinless NP536786.1
Anopheles gambiae Crossveinless/Tsg XP310820.4
Tribolium castaneum Crossveinless/Tsg XP 001812776.1
Gryllus bimaculatus Crossveinless/Tsg1 TRINITY DN44801 c8 g1 i1
Gryllus bimaculatus Crossveinless/Tsg2 TRINITY DN36254 c5 g1 i3
Oncopeltus fasciatus Crossveinless/Tsg PTY16220.1
Nasonia vitripennis Crossveinless/Tsg XP 001600362.1
Branchiostoma floridae Tsg XP002593913.1
Homo sapiens Tsg NP065699.1
Danio rerio Tsg1A NP571872.1
Homo sapiens IGFBP1 NP 000587.1
0.3
Tolloid
Homo sapiens Tolloid-like protein 2 NP036597.1
Mus musculus Tolloid-like protein 2 NP036034.1
Gallus gallus Tolloid-like 2 XP003641587.1
Homo sapiens BMP1 NP006120.1
Danio rerio BMP1a NP001035126.1
Danio rerio BMP1b NP001034901.1
Homo sapiens Tolloid-like protein 1 NP036596.3
Gallus gallus Tolloid-like protein 1 NP990034.1
Danio rerio Tolloid-like protein 1 NP571085.1
Branchiostoma floridae BMP1 AAX84844.1
Lottia gigantea Tolloid 236092
Tribolium castaneum Tolloid XP970162.1
Oncopeltus fasciatus PTY11378.1 Tolloid
Gryllus bimaculatus Tolloid TRINITY DN35005 c2 g1 i1
Nasonia vitripennis Tolloid XP 008211528.1
Drosophila melanogaster Tolloid AAF56329.2
Drosophila melanogaster Tolkin AAF56328.1
Nematostella vectensis Tolloid EDO41783.1
0.2
Gryllus
Drosophila
Tribolium
Nasonia
Oncopeltus

**Figure S6.** Crossveinless2/BMPER and KCP (Cv2), Twisted Gastrulation/Crossveinless (Tsg) and Tolloid interrelationships across the Metazoa, as determined by Bayesian (Huelsenbeck and Ronquist 2001) methods.
Crossveinless2/BMPER tree based on 213 informative residue global alignment made in MAFFT –add (Katoh and Standley 2013) with the G-iNS-i strategy with all gaps removed. Phylogeny determined with the Tree rooted with *Mus musculus* Kcp protein after (Ikeya et al 2006). Crossveinless 2/BMPER tree determined using WAG (Whelan and Goldman 2001) model. Tsg phylogeny based on a 140 informative amino acid global alignment generated using MAFFT -add (Katoh and Standley 2013) against the previously established dataset used in Kenny et al 2015 and homologues of known identity from NCBI's nr database using the G-iNS-i strategy, rooted with *H. sapiens* IGFBP (NP 000587.1) after Vilmos et al. (2001), with Tsg phylogeny determined using the Dayhoff model. Tolloid phylogeny generated according to the WAG model from a 160 informative amino acid alignment spanning the calcium-binding EGF domain and immediately proceeding the Cub domain, generated using MAFFT -add (Katoh and Standley 2013) under the G-INS-i strategy and rooted with Tolloid-like protein sequence. Scale bars represent substitutions per site at given distances. The sequences of *Gryllus bimaculatus*, *Drosophila melanogaster*, *Tribolium castaneum*, *Nasonia vitripennis* and *Oncopeltus fasciatus* are highlighted by indicated colours.

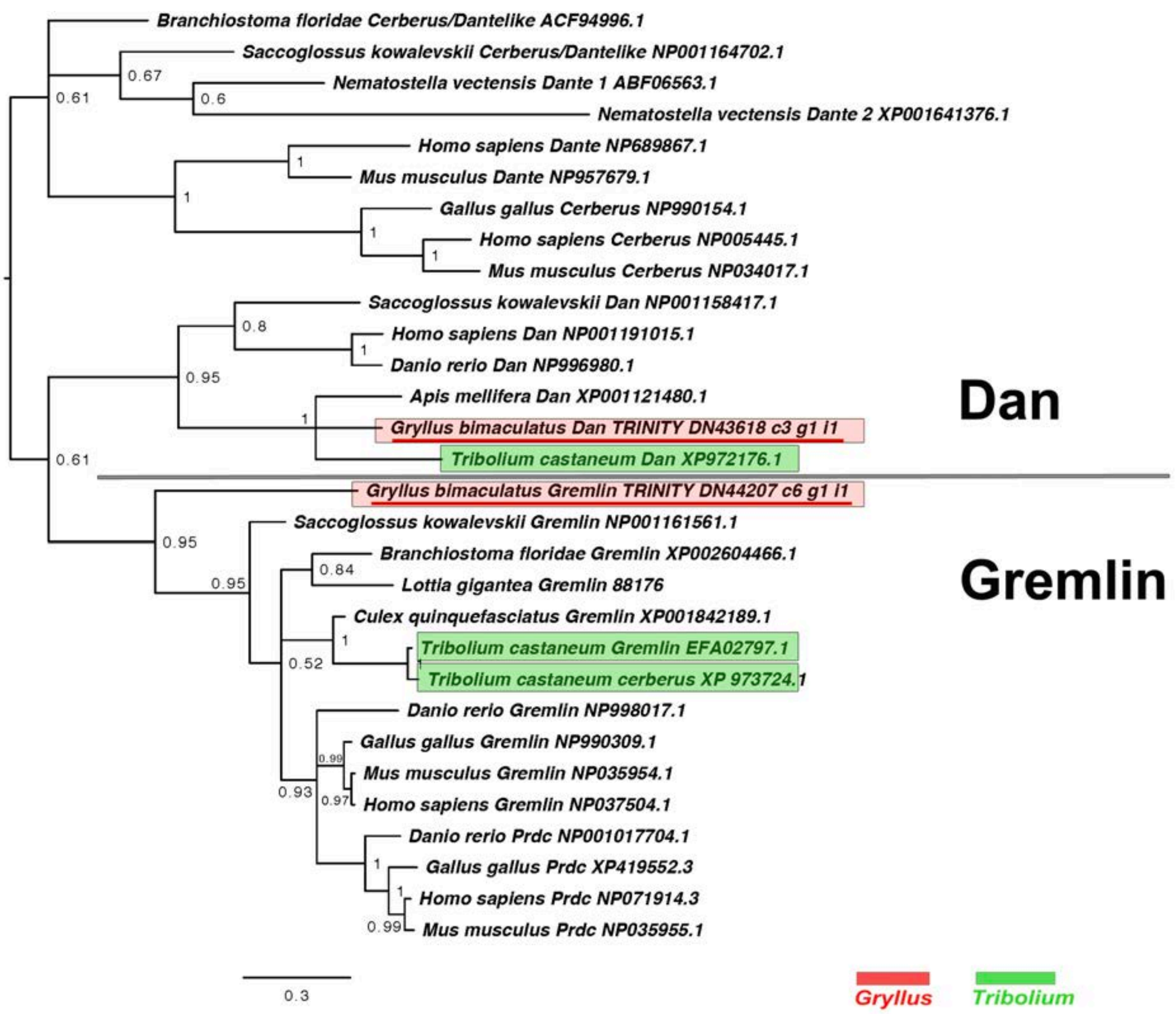

**Figure S7.** DAN class interrelationships across the Metazoa, as determined by Bayesian (Huelsenbeck and Ronquist 2001) methods.
Alignment generated using MAFFT –add (Katoh and Standley 2013) against the previously established dataset used in Kenny et al 2015 and homologues of known identity from NCBI's nr database using the G-iNS-i strategy. Phylogenies calculated on the basis of a 76 informative amino acid alignment spanning the DAN domain (Pfam ID PF03045). Phylogenies determined using the WAG model (Whelan and Goldman 2001) and rooted at midpoint. Posterior probabilities can be seen at the base of nodes. Scale bars represent substitutions per site at given distances. The sequences of *Gryllus bimaculatus and Tribolium castaneum* are highlighted by indicated colours.

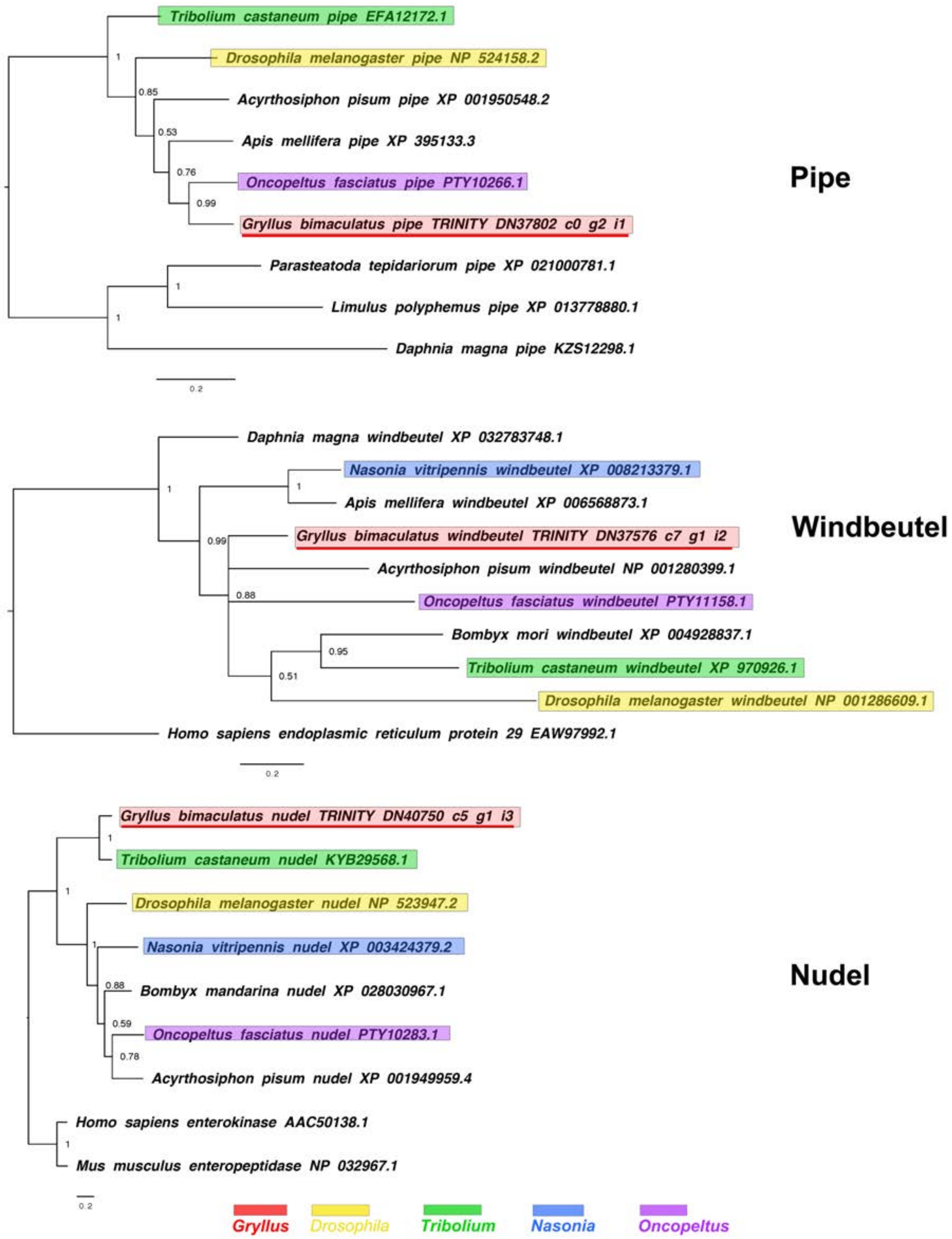


**Figure S8.** *pipe*, *windbeutel* and *nudel* interrelationships across the Metazoa, as determined by Bayesian (Huelsenbeck and Ronquist 2001) methods. *pipe* phylogeny based on a 259 informative amino acid global alignment generated using MAFFT (Katoh and Standley 2013) alongside homologues of known identity from NCBI's nr database using the G-iNS-i strategy. Tree was run for 1 million generations under the WAG model, and rooted with non-insect *pipe* sequences. *windbeutel* phylogeny based on a 227 informative amino acid global alignment generated using MAFFT (Katoh and Standley 2013) alongside homologues of known identity from NCBI's nr database using the G-iNS-i strategy, and rooted with the human outgroup sequence. Tree was run for 1 million generations under the WAG model.

*nudel* phylogenies inferred on the basis of a MAFFT alignment (Katoh and Standley 2013, G-INS-i strategy vs homologues of known identity from NCBI's nr database) with 305 informative sites, analysed under the Blosum model for 1 million generations, and shown rooted with outgroup enterokinases. *Gryllus bimaculatus* sequences underlined in red. Scale bars represent substitutions per site at given distances.

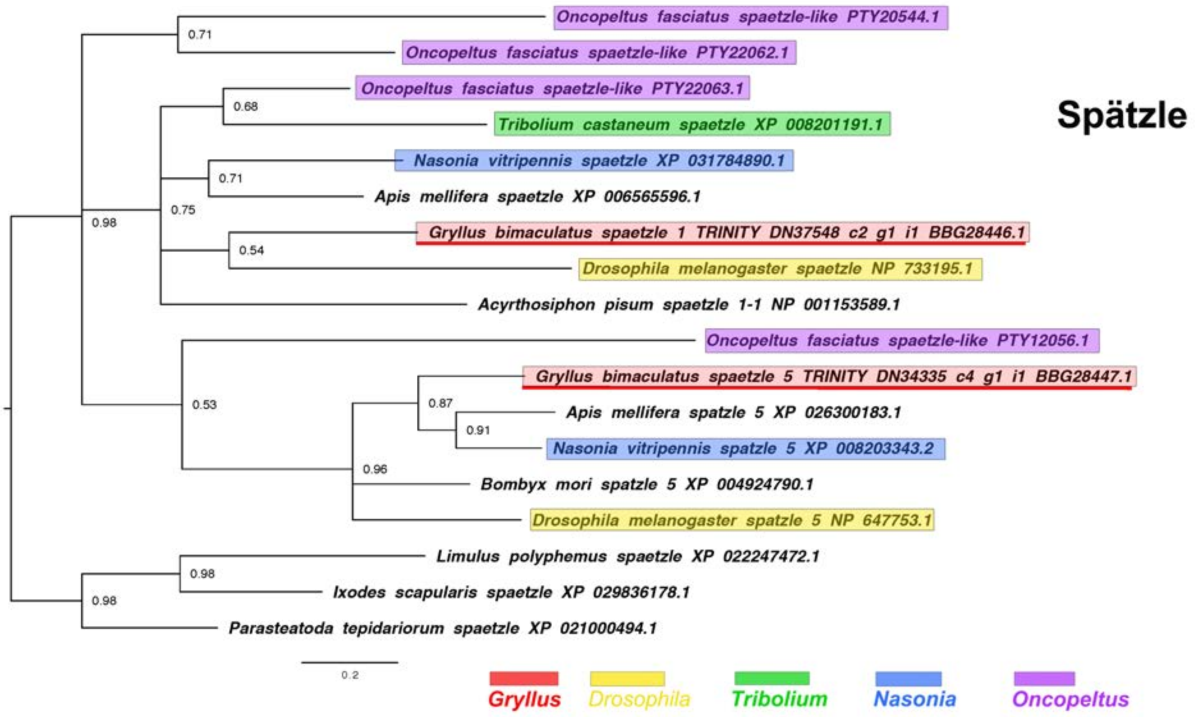


**Figure S9.** *spaetzle* interrelationships across the Metazoa, as determined by Bayesian (Huelsenbeck and Ronquist 2001) methods.
*spaetzle* phylogeny based on a 92 informative amino acid global alignment generated using MAFFT (Katoh and Standley 2013) alongside homologues of known identity from NCBI's nr database using the G-iNS-i strategy, analysed under the WAG model for 2 million generations, and rooted with non-insect *spaetzle* sequences. NB in some *Oncopeltus* sequences excluded due to truncated sequence lying outside domain used for phylogenetic reconstruction. *Gryllus bimaculatus* sequences underlined in red. Scale bars represent substitutions per site at given distances.

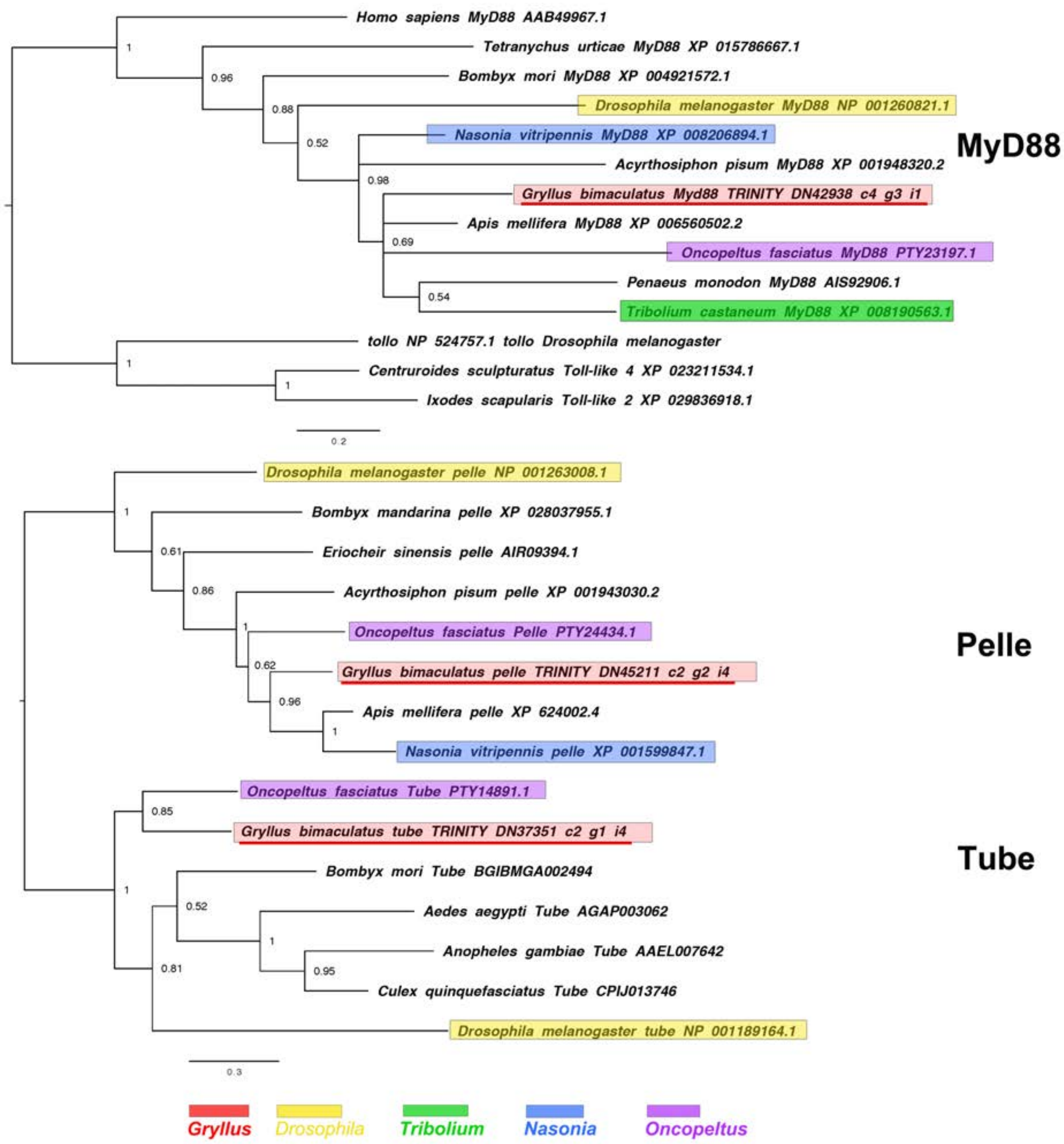


**Figure S10.** *pelle* and *tube* interrelationships across the Metazoa, as determined by Bayesian (Huelsenbeck and Ronquist 2001) methods. *pelle* and *tube* phylogenies inferred on the basis of a MAFFT alignment (Katoh and Standley 2013, G-INS-i strategy vs homologues of known identity from NCBI's nr database) with 132 informative sites, and shown with the root chosen between the two gene families. Tree was run for one million generations under the WAG model. *Gryllus bimaculatus* sequences underlined in red. Scale bars represent substitutions per site at given distances. *MyD88* phylogeny based on a 126 informative amino acid global alignment generated using MAFFT L-INS-I (Katoh and Standley 2013) alongside homologues of known identity from NCBI's nr database. Tree was run for 1 million generations under the WAG model, rooted with Toll-like outgroups. *Gryllus bimaculatus* sequences underlined in red. Scale bars represent substitutions per site at given distances.

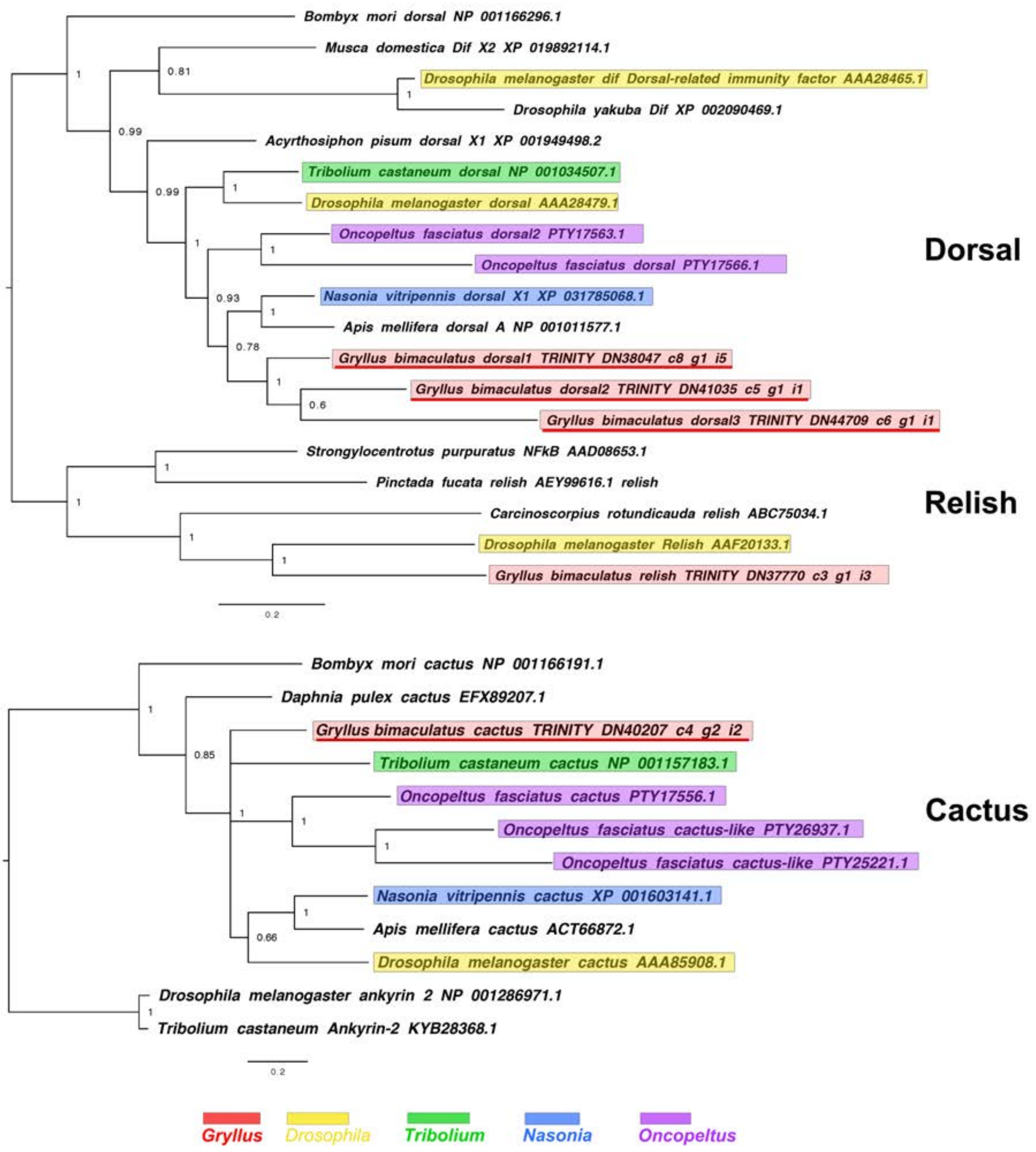


**Figure S11.** *dorsal*, *relish* and *cactus* interrelationships across the Metazoa, as determined by Bayesian (Huelsenbeck and Ronquist 2001) methods. *dorsal* phylogeny based on a 221 informative amino acid global alignment generated using MAFFT (Katoh and Standley 2013) alongside homologues of known identity from NCBI's nr database using the G-iNS-i strategy, analysed under the WAG model for 2 million generations, and rooted with *relish* sequences (including that of *Gryllus bimaculatus)*. *cactus* phylogenies inferred on the basis of a MAFFT alignment (Katoh and Standley 2013, G-INS-i strategy vs homologues of known identity from NCBI's nr database) with 122 informative sites, analysed under the cprev model for 1 million generations, and shown rooted with ankyrin 2 sequences. Scale bars represent substitutions per site at given distances. NB for *cactus* some *Oncopeltus* sequences excluded due to truncated sequence lying outside domain used for phylogenetic reconstruction.

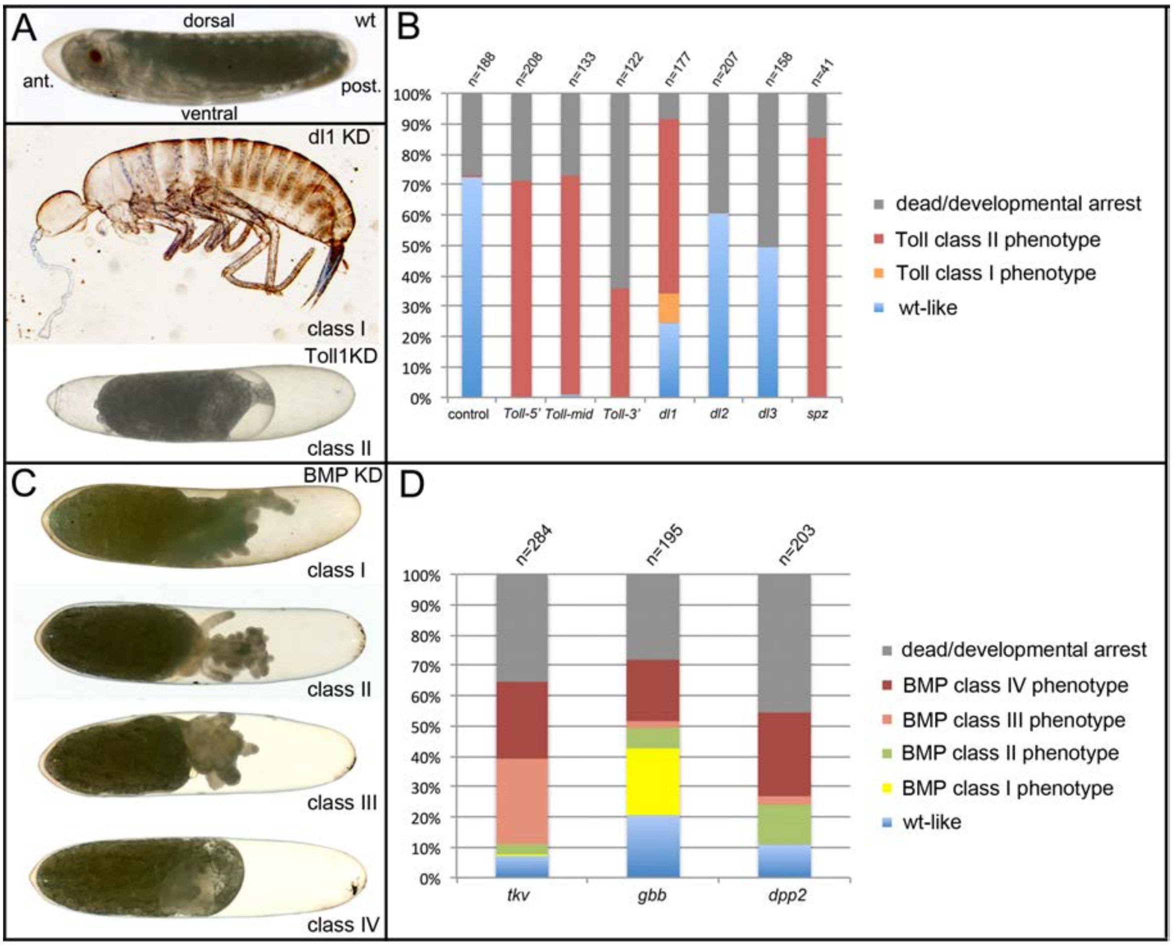


**Figure S12.** Phenotype classes resulting from reduced Toll and BMP signalling.
**(A)** Control wildtype (wt) embryo and terminal phenotypes produced by KD of Toll signalling components. The wildtype larva is at egg stage 20. The anterior pole of the egg is pointed, the dorsal side concave, the ventral side convex. Weak (class I) phenotypes (*Gb-dl1* KD) are characterized by the production of fully segmented larvae, which completed katatrepsis and dorsal closure. Patterning defects are strongest in the thoracic and head region with cuticle constrictions, eye fusions or deletion of eyes (see also Figure S17). Strong (class II) phenotypes (here *Gb-Toll1* KD) do not produce recognizable cuticle structures. Embryonic tissue fragments remain engulfed within the yolk (Figure S16). The serosa contracts from both anterior and posterior poles leaving large fluid-filled spaces at both egg poles (see also Figure S21 and video S2). **(B)** Frequency of phenotypic classes upon pRNAi of Toll signalling components (three non-overlapping fragments of *Gb-Toll1*, *Gb-dl1*, *Gb-dl2*, *Gb-dl3* and *Gb-spz*). Weak phenotypes (Class I) were rare and only found upon *Gb-dl1* KD. **(C)** Terminal phenotypes produced by KD of BMP signalling components. Weak (class I) phenotypes possess thoracic and head appendages. Eyes are frequently present, but may be fused at the dorsal side indicating a loss of dorsal tissue and compensating expansion of ventral structures (Figure S17). The abdomen is reduced to a tube-like structure (Figure S22). Medium strong (class II) phenotypes carry only head segments, most prominently the antennae. Like in class I the abdomen is reduced to a tube-like structure (Figure S22). Strong (class III) phenotypes lack recognizable segmental structures altogether. Class IV embryos have not undergone katatrepsis and thus remain engulfed within yolk and serosa. As this phenotype occurs independent from degree of segmental defects used to distinguish class I, class II and class III, its strength cannot be determined. **(D)** Frequency of phenotypic classes upon pRNAi of BMP signalling components.

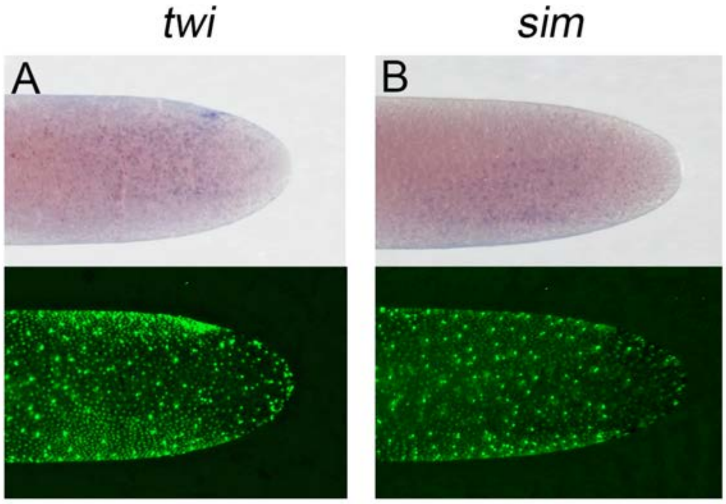


**Figure S13.** Expression of *Gb-twi* and *Gb-sim* in early *Gryllus* embryos.
Whole mount ISH for indicated genes and DNA staining (Sytox) showing ventral surface views of the posterior 40% of embryos at early germ anlage condensation (ES 2.2-2.3).

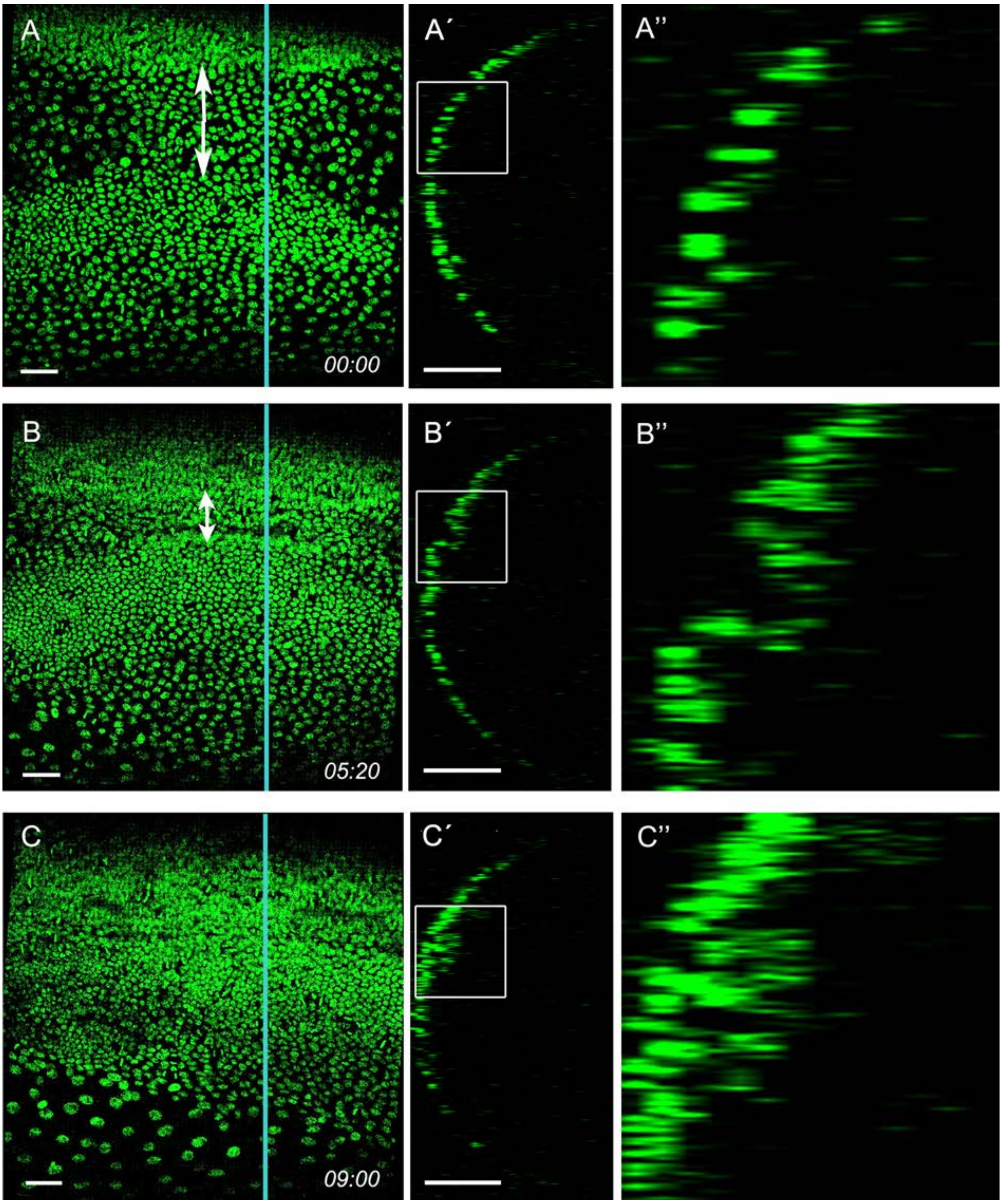


**Figure S14.** Mesoderm internalisation.
Stills from video 1. Ventral surface view and z-sections of embryo carrying a pXLBGact Histone2B:eGFP transgene (A) Early germ anlagen condensation (ES 2.3). (B) Mid germ anlagen condensation (ES 2.4-2.5). (C) Late germ anlagen condensation (ES 2.6). The z-sections show that cells positioned between the lateral plates of high cell density become internalized as the plates move ventrally. Staging according to (Donoughe and Extavour, 2016; Sarashina et al., 2005).

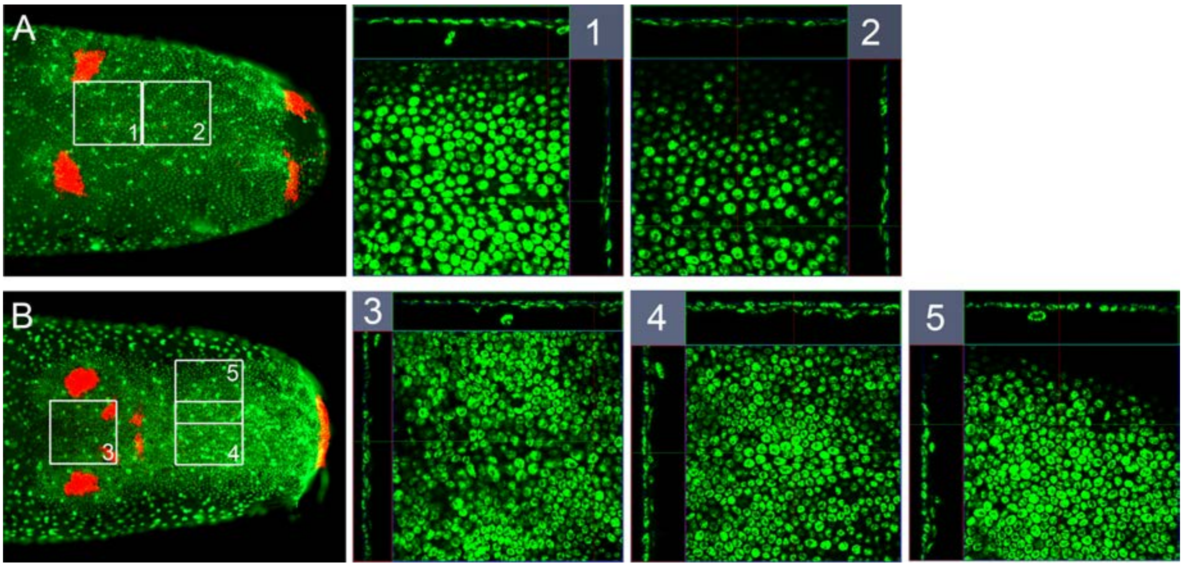


**Figure S15.** Mesoderm in early germ band embryos.
Whole mount ISH for *Gb-wg* and DNA staining (Sytox) showing ventral surface views of the posterior 30% of embryos and z-sections at indicated positions **(A)** shortly after ventral fusion of the lateral plates (early ES 3,0, only ocular and posterior *Gb-wg* domains) and **(B)** at early germ band elongation (late ES 3.0, additional antennal and mandibular *Gb-wg* stripes). In **(A)** a thin inner cell layer is visible in central regions of the germ band. In **(B)** the inner cell layer becomes more pronounced. Staging according to (Donoughe and Extavour, 2016; Sarashina et al., 2005).

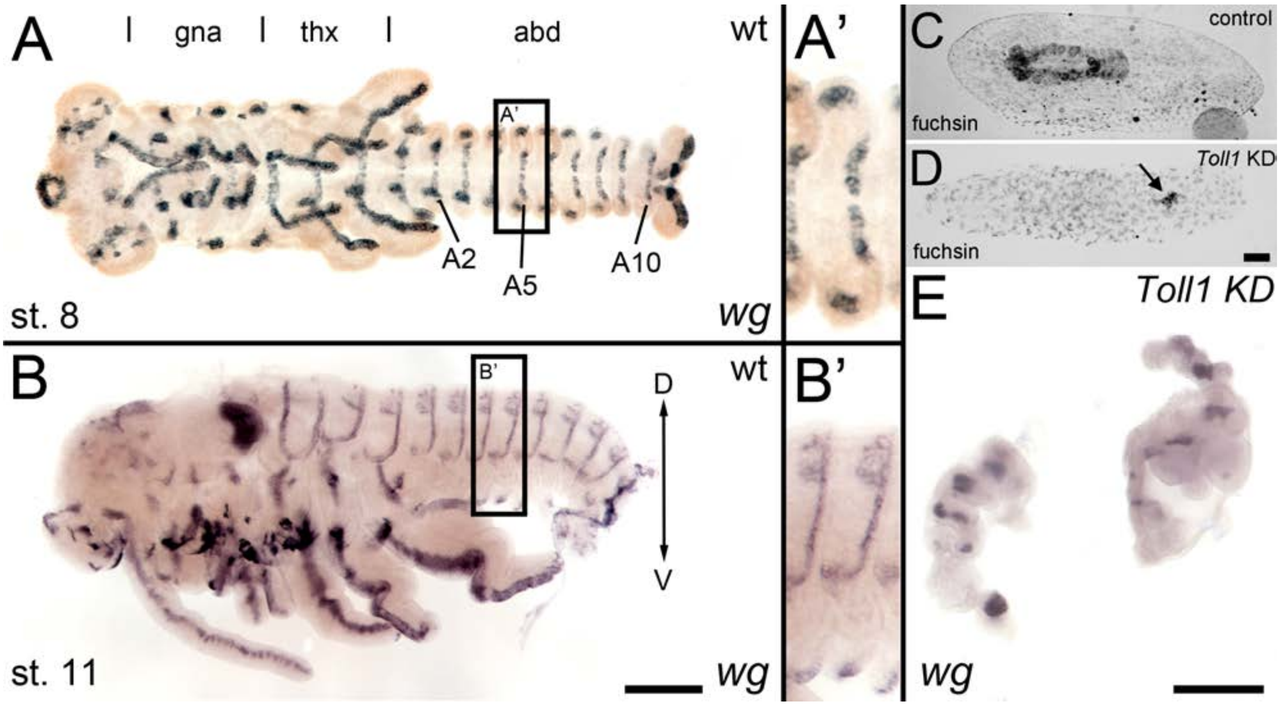


**Figure S16.** *wingless* expression in late *Gb-Toll1* KD embryos.
*Gb-wg* expression in control embryos and *Gb-Toll1* KD embryos. **(A)** Ventral view of ES 8 embryo. Different tagmata (gna, gnathal; thx, thorax; abd, abdomen) and abdominal segments 2, 5 and 10 (A2, A5, A10) are indicated. **(A')** *Gb-wg* stripes are largely restricted to the ventral ectoderm. **(B)** Lateral view of ES 11 embryo. **(B')** At this stage *Gb-wg* becomes expressed in the dorsal ectoderm. **(C)** Fuchsin staining of control embryo after anatrepsis. The embryo has been internalized into the yolk. **(D)** Anatrepsis also occurs in *Gb-Toll1* KD embryos (see video 5), however the embryo remains engulfed within the yolk. It consists of small tissue fragments (arrow), which express *Gb-wg* **(E)**. As *Gb-Toll1* KD embryos lack segmental *Gb-wg* expression prior to anatrepsis (Figure 8) we assume that the *Gb-wg* expression in late embryos corresponds to the dorsal ectodermal *Gb-wg* stripes of control embryos. Staging according to (Donoughe and Extavour, 2016; Sarashina et al., 2005).

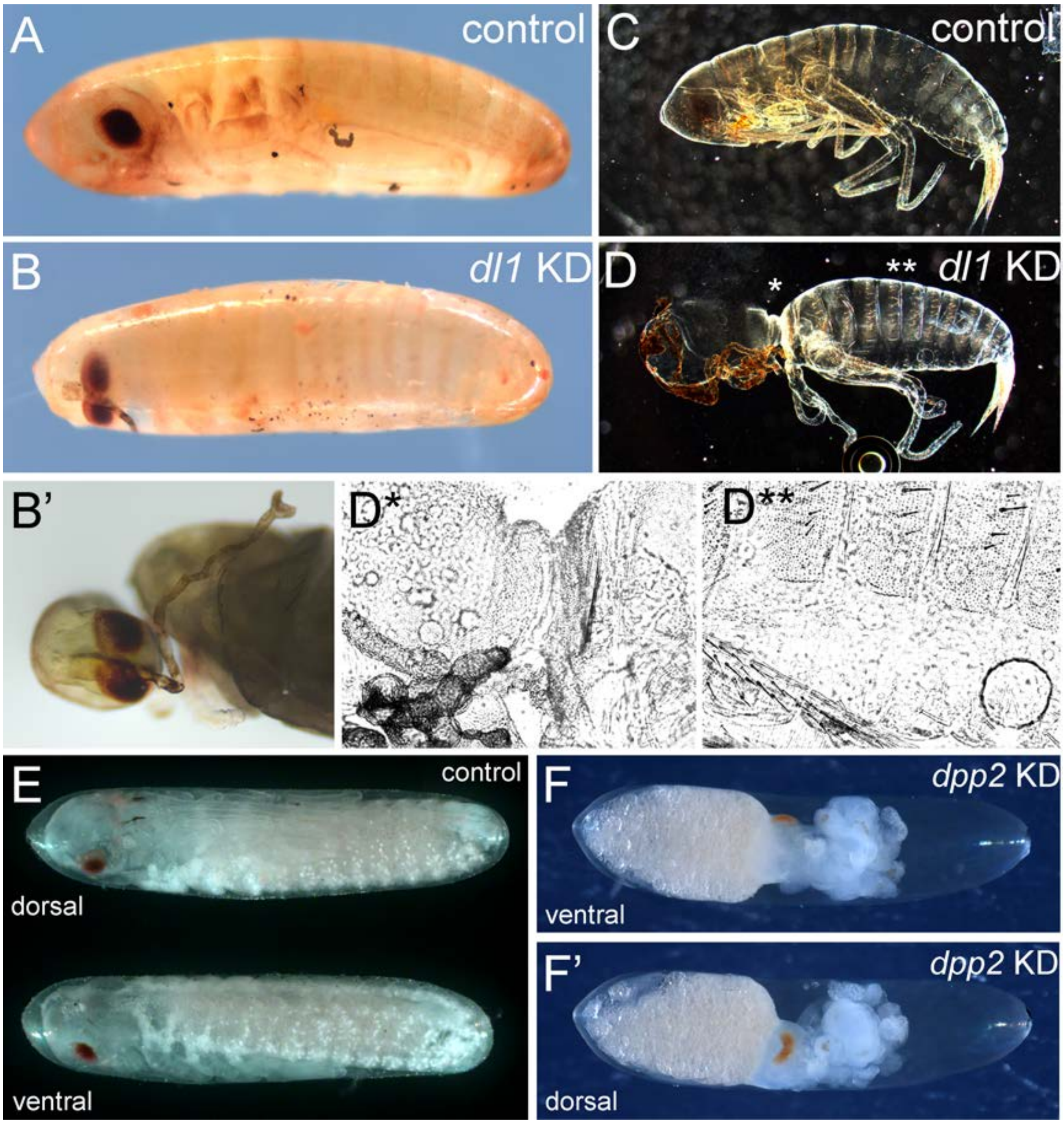


**Figure S17.** Later phenotypes produced by interfering with Toll and BMP signalling. **(A, B)** Bright field micrograph of control **(A)** and *Gb-dl1* KD **(B, B')** embryo at egg stage 22. The ventral fusion of eyes and probably also of the antennae indicates a weak dorsalisation (i.e. a loss of ventral tissue compensated by expansion of dorsal tissue). **(C, D)** Dark field micrograph of control **(C)** and *Gb-dl1* KD **(D)** larvae after completion of embryonic development. **(D*, D**)** Phase contrast micrographs of positions indicated in **(D)** reveal that the constricted region is composed of cuticle with dorsal identity. **(E)** Ventral and dorsal views of control embryo at egg stage 20. **(F, F')** Ventral and dorsal views of *Gb-dpp2* KD. The presence of anterior appendages (Figure S22) demonstrates ventrolateral tissue specification. Therefore, we assume that the eyes are fused at the dorsal side indicating a ventralisation (i.e. loss of dorsal tissue compensated by expansion of ventral tissue). Staging according to (Donoughe and Extavour, 2016)

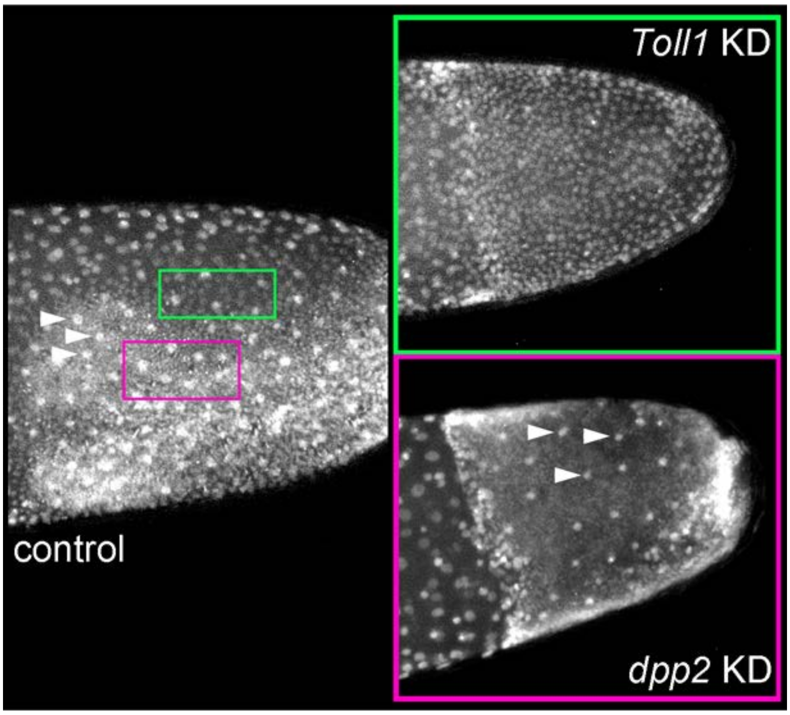


**Figure S18.** Cell densities in embryos lacking Toll or BMP signalling.
Stills from videos of pXLBGact Histone2B:eGFP embryos. Control and KD embryos are shown at corresponding stages (early germband stage approximately ES 3.0). Regions of low (green) and high (pink) cell densities are demarcated in the control embryo. Arrowheads indicate yolk nuclei. The cell density of the germband of *Gb-Toll1* KD embryos corresponds to low cell density regions found at the dorsal margin of the germband in control embryos (green). The cell density of the outer layer of the germband of *Gb-dpp2* KD embryos corresponds to high cell density regions found in ventral parts of germband in control embryos (pink). Staging according to (Donoughe and Extavour, 2016; Sarashina et al., 2005).

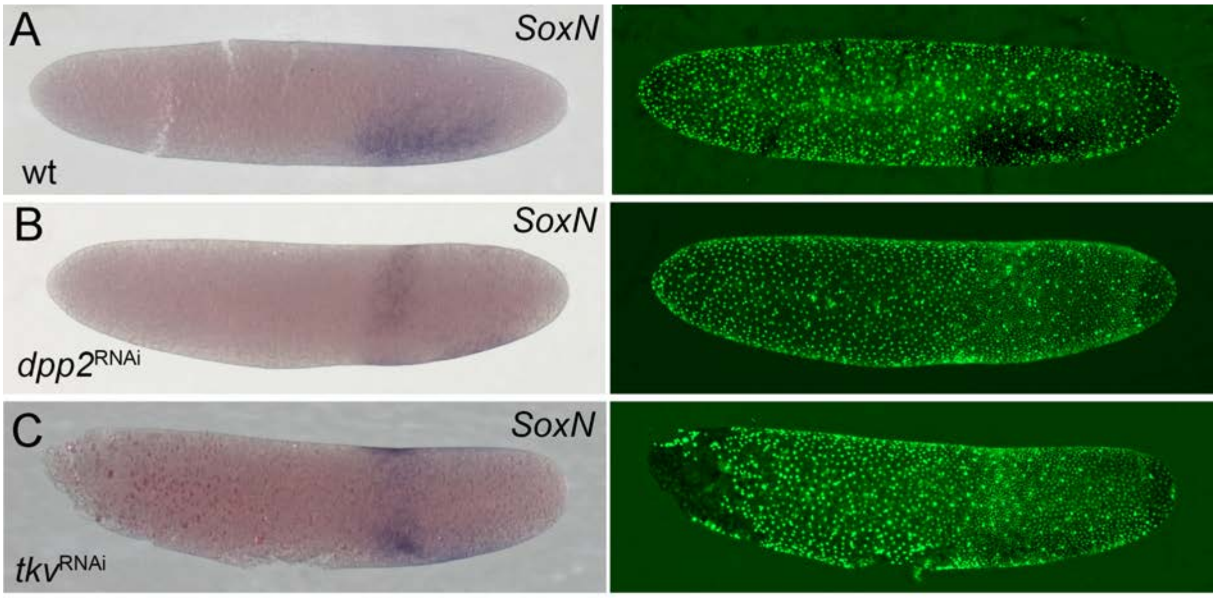


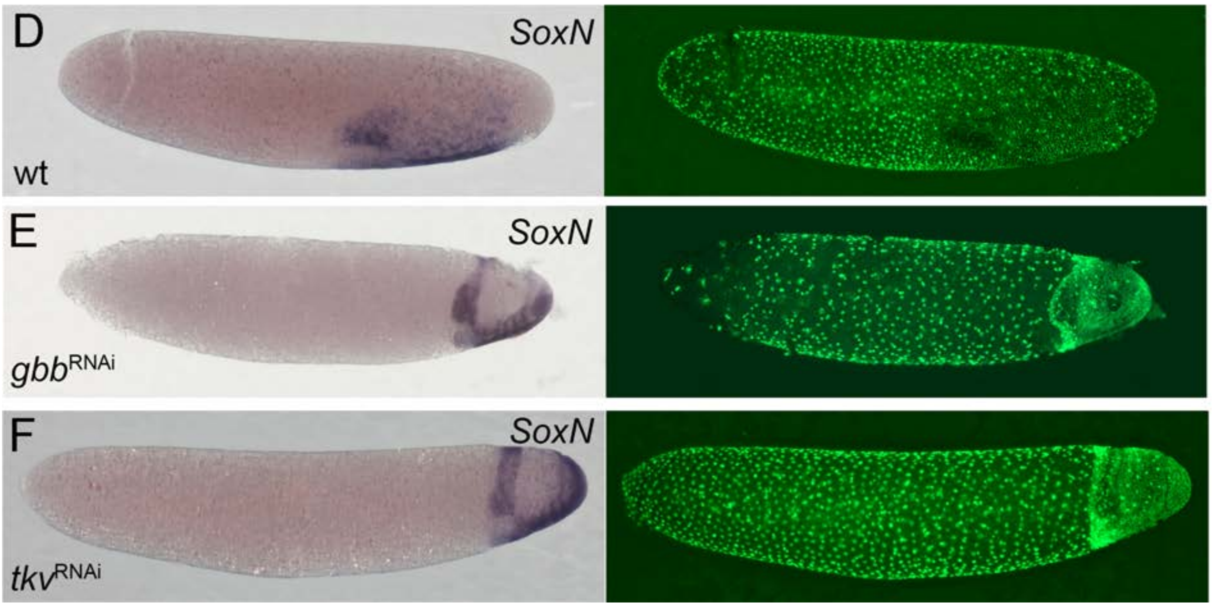


**Figure S19.** Comparison of DV fate map shift after *Gb-tkv*, *Gb-dpp2* and *Gb-gbb* KD. Whole mount ISH (purple) for *Gb-SoxN* and DNA staining (Sytox, green). **(A-C)** Early germ anlage condensation (ES 2.2-2.3). **(D-F)** Late germ anlage condensation to early germband (ES 2.4 -3.0). Staging according to (Donoughe and Extavour, 2016; Sarashina et al., 2005)

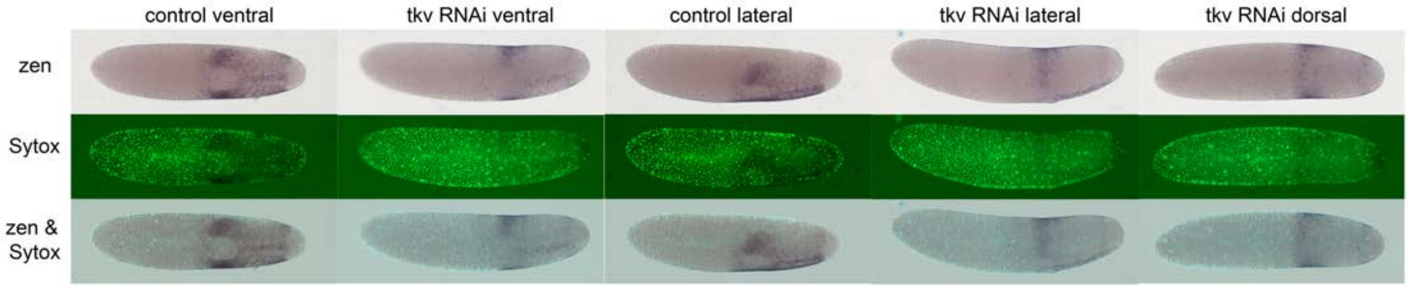


**Figure S20.** *Gb-zen* expression upon *Gb-tkv* KD.
Whole mount ISH (purple) for *Gb-zen*, DNA staining (Sytox, green) and overlay of control and *Gb-tkv* KD embryos at late germ anlage condensation (ES 2.4-2.6). Ventral surface views show that the ventral expression domain of *Gb-zen* is not expanded upon *Gb-tkv* KD. In contrary the head expression domain expands evenly to the dorsal side as seen by lateral and dorsal surface views. Staging according to (Donoughe and Extavour, 2016; Sarashina et al., 2005).

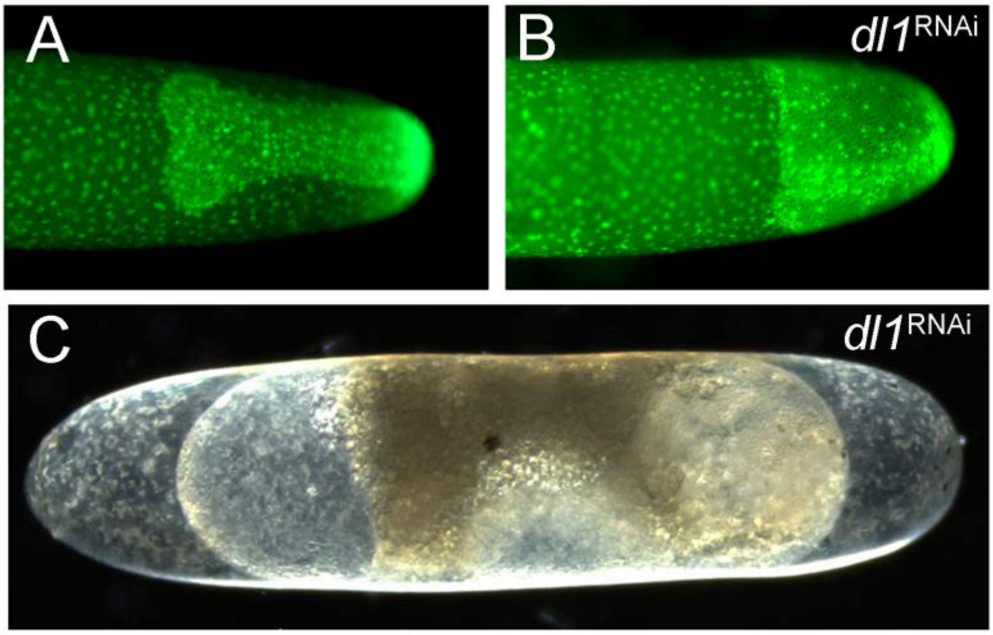


**Figure S21.** The strong phenotype of *Gb-dl1* KD.
(A, B) Stills from videos of pXLBGact Histone2B:eGFP embryos. (A) Control embryo at early germband elongation. (B) *GB-dl1* KD embryo at corresponding stage. The embryo is rotationally symmetric lacking DV polarity. (C) The terminally differentiated KD embryo shows the contraction of the serosa from anterior and posterior poles, which is typical for lack of Toll signalling in *Gryllus*.

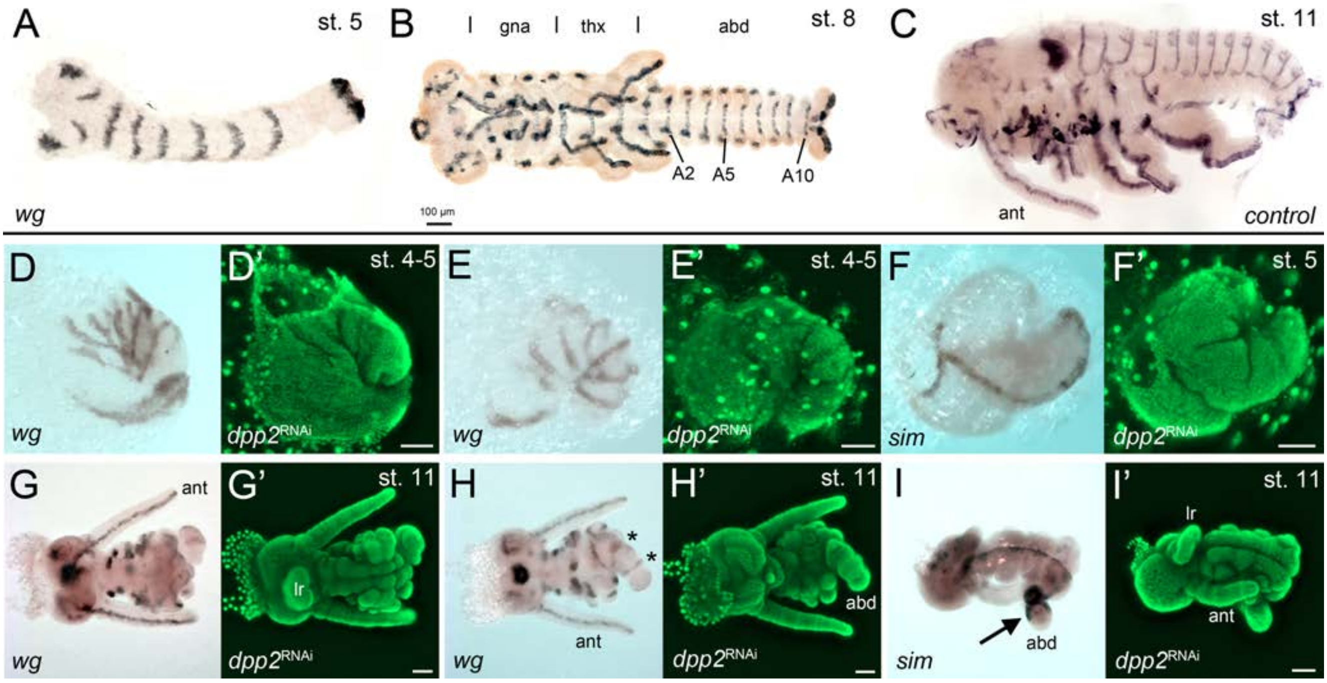


**Figure S22.** The development of *Gb-dpp2* KD embryos.
(**A-C**) *Gb-wingless* expression in control embryos at different developmental stages. The different tagmata (gna, gnathal; thx, thorax; abd, abdomen) and abdominal segments 2, 5 and 10 (A2, A5, A10) are indicated in B. (**D, E**) Posteriorly condensed *Gb-dpp2* KD embryos show irregular *Gb-wg* stripes (compare to **A**). However, seven stripes can be distinguished suggesting that segmentation occurred in the head and thoracic region. (**F**) Although the morphology of the early embryos was strongly altered, ventral midline expression of *Gb-sim* was present. (**G, H**) Ventral **(G)** and dorsal **(H)** of ES 11 KD embryo. Regular *wg* expression is visible in the antennae (ant) and the labrum (lr). The asterisk in H marks the remaining segments in the tube-like abdomen. Abdominal segment number and gnathal and thoracic appendages were strongly reduced. (**I, I'**) *Gb-sim* expression in ES 11 KD embryo. The remaining tube-like abdominal segments are strongly marked by *Gb-sim* expression (arrow in I). Staging according to (Donoughe and Extavour, 2016; Sarashina et al., 2005).

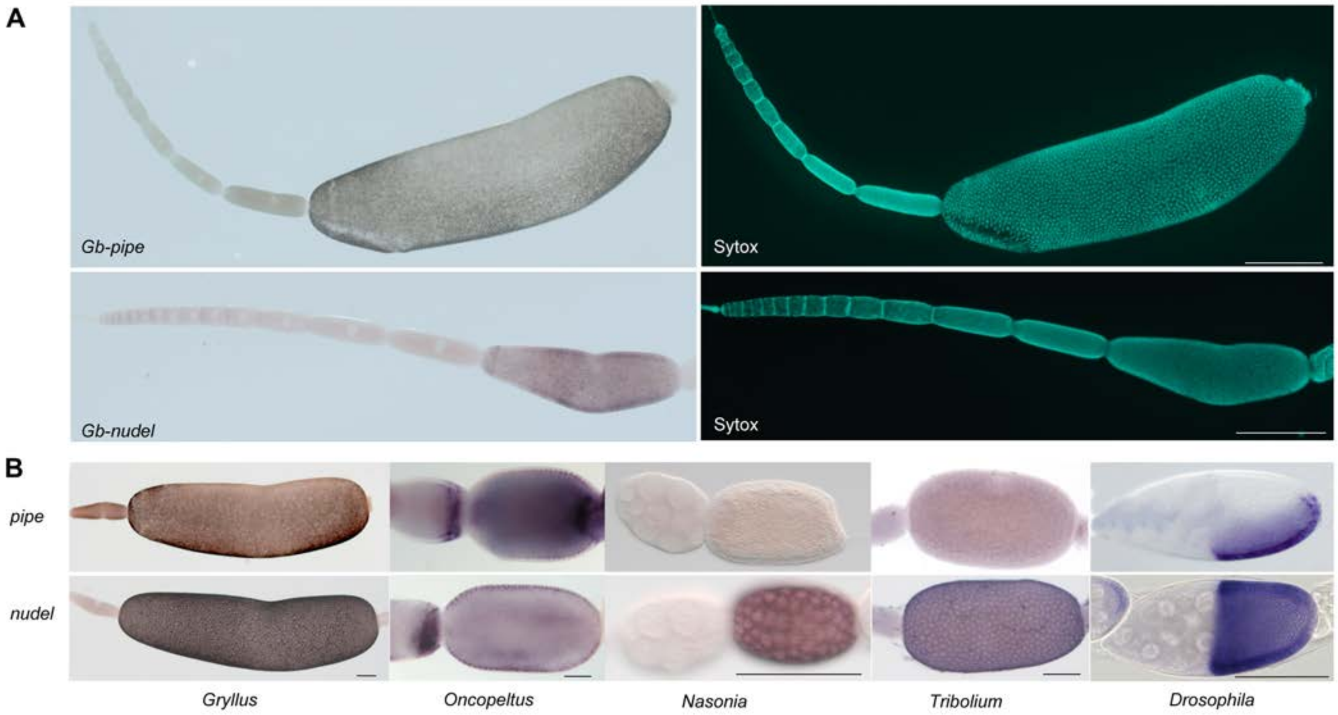


**Figure S23.** Follicle cell expression of *pip*e and *nudel* in different insects.
**(A)** *Gb-pipe* and *Gb-nudel* ISH and DNA staining (Sytox) in *Gryllus* ovarioles. The anterior tip with the terminal filament and the germarium is pointing to the left side. *Gb-nudel* and *Gb-pipe* expression are detected in the follicular epithelium surrounding the oocyte after asymmetric positioning of the oocyte nucleus inducing a shape change in the follicular epithelium (Lynch et al., 2010). The scale bars correspond to 500 μm. **(B)** *pipe* and *nud*el ISH in egg chambers of the indicated insects. While uniform *nudel* expression was detected for all studied insects (Chen, 2015; Hong and Hashimoto, 1995), *pipe* was found to be expressed in follicle cells only in *Drosophila* (Sen et al., 1998*)*, *Oncopeltus* (Chen, 2015) and *Gryllus*. A dorsal repression of *pipe* was only observed in *Drosophila* and *Gryllus*. The scale bars correspond to 500 μm in A and 100 μm in B.

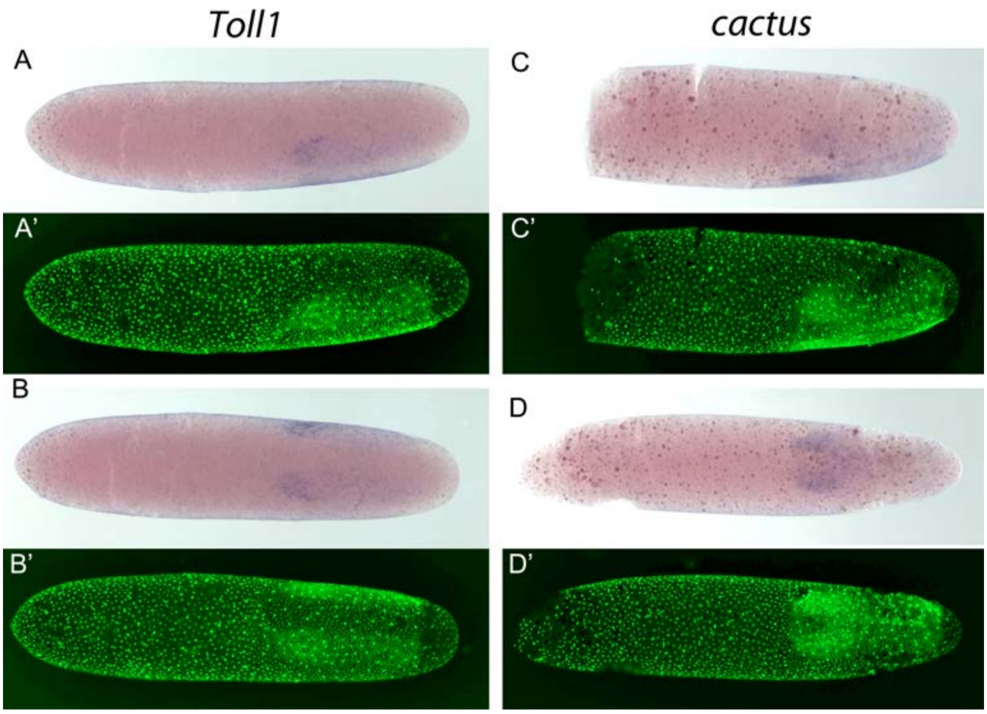


**Figure S24**. Expression of *Gb-Toll1* and *Gb-cactus*.
*Gb-Toll1* and *Gb-cactus* ISH (blue) and DNA staining (Sytox, green). **(A)** Lateral view of embryo at late germ anlage condensation. **(B)** Ventral view of the embryo shown in (A). *Gb-Toll1* is upregulated in the ectoderm with stronger expression in the head lobes. No expression in seen in the prospective mesoderm. In *Tribolium, Toll1* expression is upregulated in ventral regions including the prospective mesoderm where the nuclear Dorsal/NF-kB protein levels are high indicating positive feedback regulation between the Toll receptor and its downstream transcription factor. **(C)** Lateral view of embryo at late germ anlage condensation. **(D)** Ventral view of embryo after mesoderm internalisation. *Gb-cact* is expressed evenly in the embryonic anlage. In *Tribolium, cactus* is upregulated in the prospective mesoderm indicating negative feedback regulation as Cactus prevents the nuclear transport of Dorsal/NF-kB protein.

### Supplementary Tables

| *TGFβ Ligands* | | *Isoforms ?* | | | *Isoforms?* |
|---|---|---|---|---|---|
| *dpp* | *TRINITY_DN43272_c4_g1_i1* | | *BAMBI* | *no clear candidate* | |
| *dpp2* | *TRINITY_DN36587_c2_g1_i1* | | *Cripto* | *no clear candidate* | |
| *gbb* | *TRINITY_DN36559_c5_g1_i1* | | *Crossveinless/Tsg* | *TRINITY_DN44801_c8_g1_i1* | *2* |
| *ADMP* | *TRINITY_DN39293_c1_g1_i6* | *5* | | *TRINITY_DN36254_c5_g1_i3* | *3* |
| *BMP3 (GDF10)* | *TRINITY_DN40222_c2_g1_i2* | *2* | *Kielin/Chordin-like* | *no clear candidate* | |
| *Myostatin (D, T)* | *TRINITY_DN40630_c10_g1_i1* | | *Crossveinless 2/ BMPER* | *TRINITY_DN38054_c10_g1_i1* | |
| *Inhibin/Activin* | *TRINITY_DN38503_c6_g1_i1* | *2* | *Dan* | *TRINITY_DN43618_c3_g1_i1* | |
| *Maverick* | *TRINITY_DN40148_c2_g1_i1* | | | | |
| *BMP15/GDF9* | *TRINITY_DN64555_c0_g1_i1* | | *Gremlin* | *TRINITY_DN44207_c6_g1_i1* | |
| *ALKs* | | | | | |
| *BMP receptor1* | *TRINITY_DN40849_c4_g1_i4* | *4* | *Follistatin* | *TRINITY_DN42584_c3_g1_i1* | |
| *TGF receptor 1 (?)* | *TRINITY_DN34065_c3_g1_i3* | *2* | | | |
| *activin receptor saxophone* | *TRINITY_DN39845_c0_g1_i1* | | *Noggin* | *TRINITY_DN43642_c1_g1* | *2* |
| *activin receptor 2* | *TRINITY_DN35393_c5_g2_i2* | *2* | | | |
| *BMP receptor 2* | *TRINITY_DN39452_c4_g1_i1* | *2* | *Short gastrulation /Chordin* | *no clear candidate* | |
| *SMADs* | | | | | |
| *MAD* | *TRINITY_DN44474_c5_g1* | *4* | *SMURF* | *TRINITY_DN37982_c4_g1_i1* | |
| *DAD* | *TRINITY_DN43199_c12_g1_i1* | | | | |
| *Medea* | *TRINITY_DN39054_c2_g4* | *2* | *Tolloid* | *TRINITY_DN35005_c2_g1* | *2* |
| *Smox* | *TRINITY_DN36276_c1_g2_i7* | *3* | | | |
| | | | *Pentagone:* | *TRINITY_DN35624_c3_g2_i5* | *10* |
| *Brinker* | *TRINITY_DN38911_c1_g1_i1* | | | | |
| | | | *Schnurri:* | *TRINITY_DN35805_c2_g1_i1* | |

**Table S1**. Recovery of the TGFβ/BMP pathway of *G. bimaculatus,* Summary. Where multiple isoforms are present, this is noted in the appropriate column.

| | |
|---|---|
| *pipe* | *TRINITY_DN37802_c0_g2_i1* |
| *windbeutel* | *TRINITY_DN37576_c7_g1_i2* |
| *nudel* | *TRINITY_DN40750_c5_g1_i3* |
| *spätzle 1-like* | *TRINITY_DN37548_c2_g1_i1* |
| *spätzle 5-like* | *TRINITY_DN34335_c4_g1_i1* |
| *Toll1* | *TRINITY_DN36150_c1_g2_i4* |
| *Myd88* | *TRINITY_DN42938_c4_g3_i1* |
| *tube* | *TRINITY_DN37351_c2_g1_i4* |
| *pelle* | *TRINITY_DN45211_c2_g2_i4* |
| *cactus* | *TRINITY_DN40207_c4_g2_i2* |
| *dorsal 1* | *TRINITY_DN38047_c8_g1_i5* |
| *dorsal 2* | *TRINITY_DN41035_c5_g1_i1* |
| *dorsal 3* | *TRINITY_DN44709_c6_g1_i1* |

**Table S2.** Recovery of Toll pathway components of *G. bimaculatus*

#### Supplementary Files

**File S1.** Data for HMM-based identification of Sog fragments in Orthoptera

**File S2.** Diamond BlastX annotations of transcriptome, by best-hit sequence in the *nr* database. 101,760 of the 328,616 contigs were annotated with an $E$ value cutoff of $10^{-6}$. Gzipped (.gz) text file.

**File S3.** Sequences, alignments (raw and trimmed), unedited tree files and tree figures in a variety of formats. Zipped (.zip) directory.
