## Supplementary figures and images for "Striking parallels between dorsoventral patterning in *Drosophila* and *Gryllus* reveal a complex evolutionary history behind a model gene regulatory network"

### ALKs.pdf

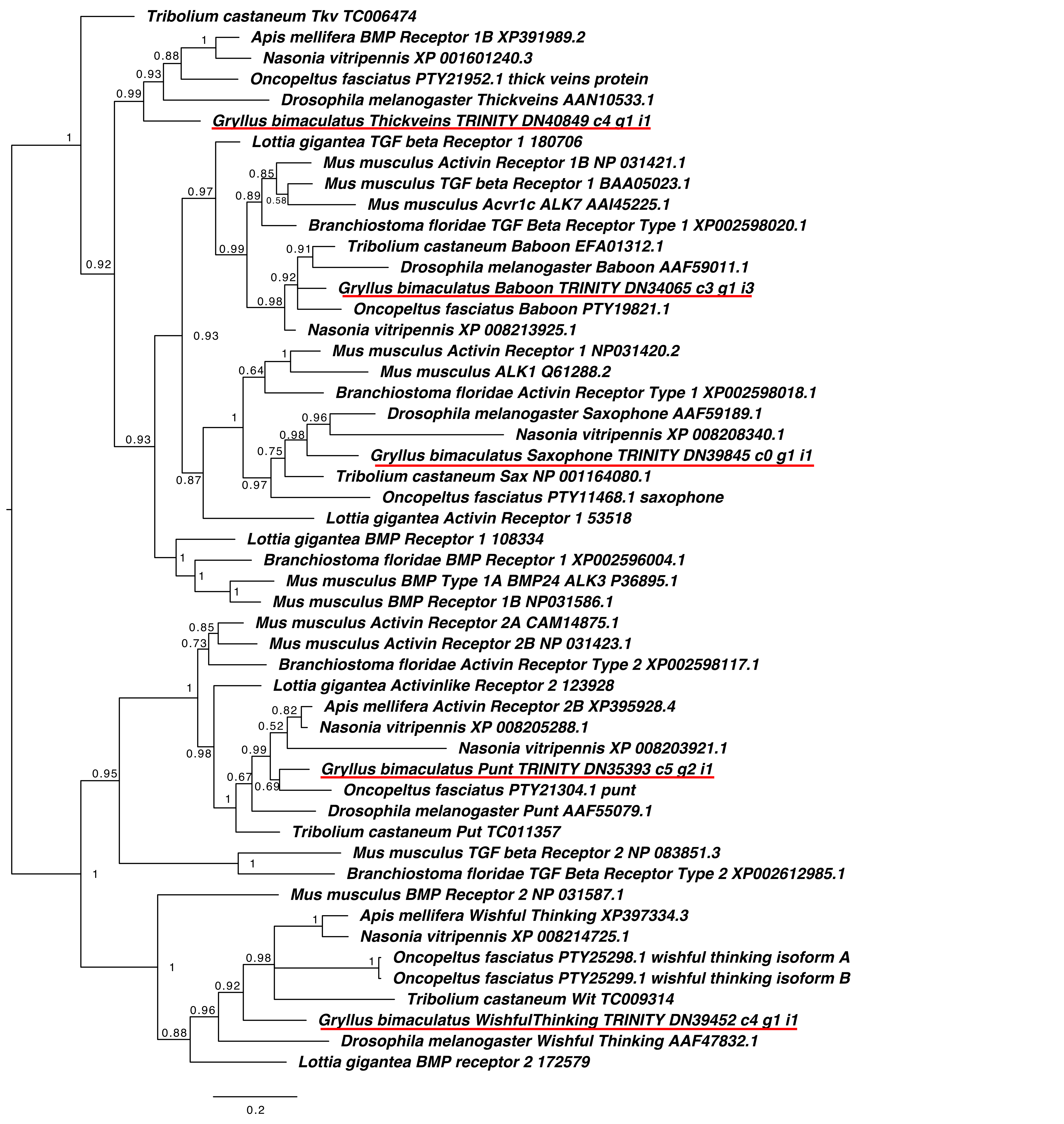

### brinker.pdf

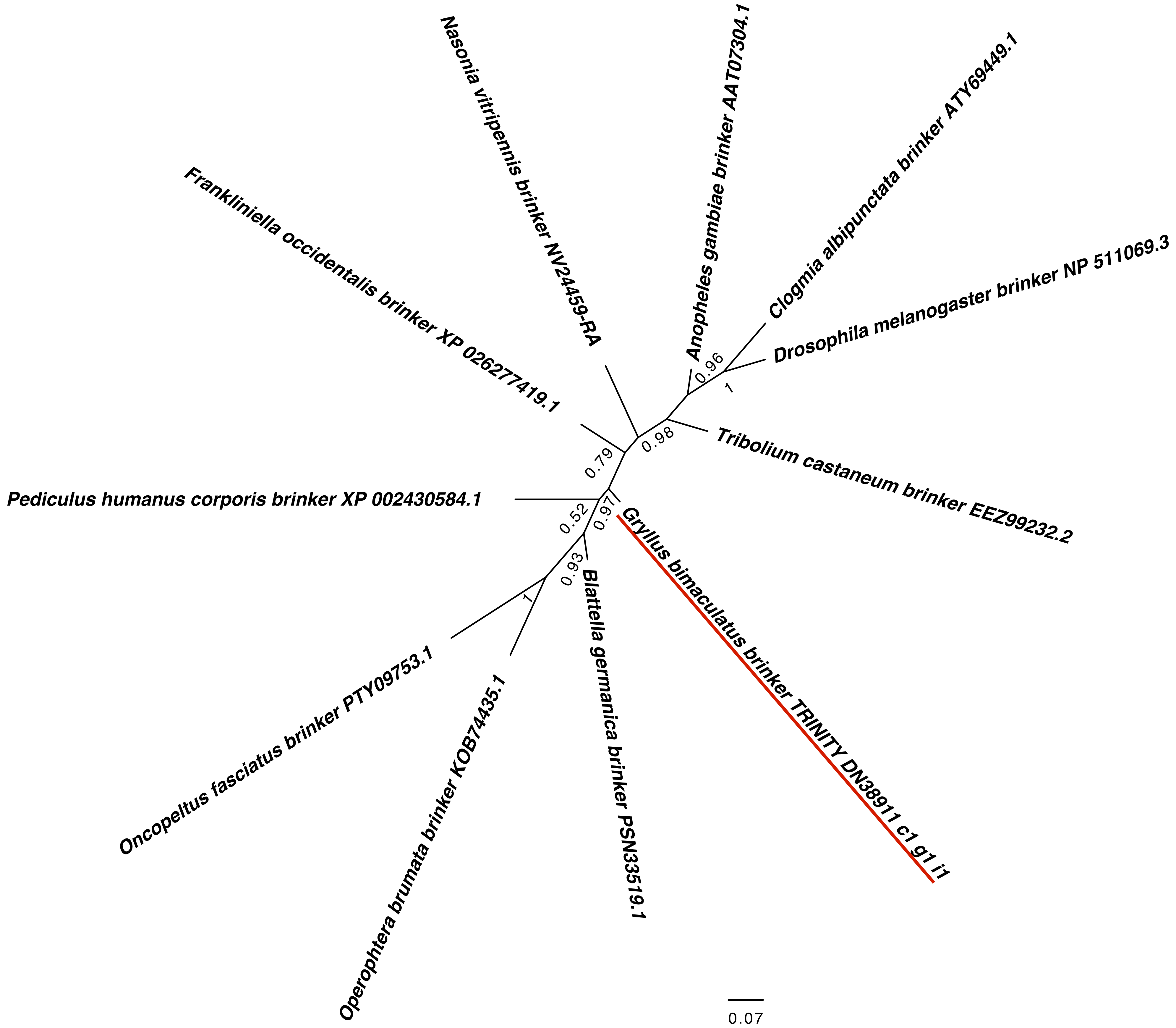

### cactus.pdf

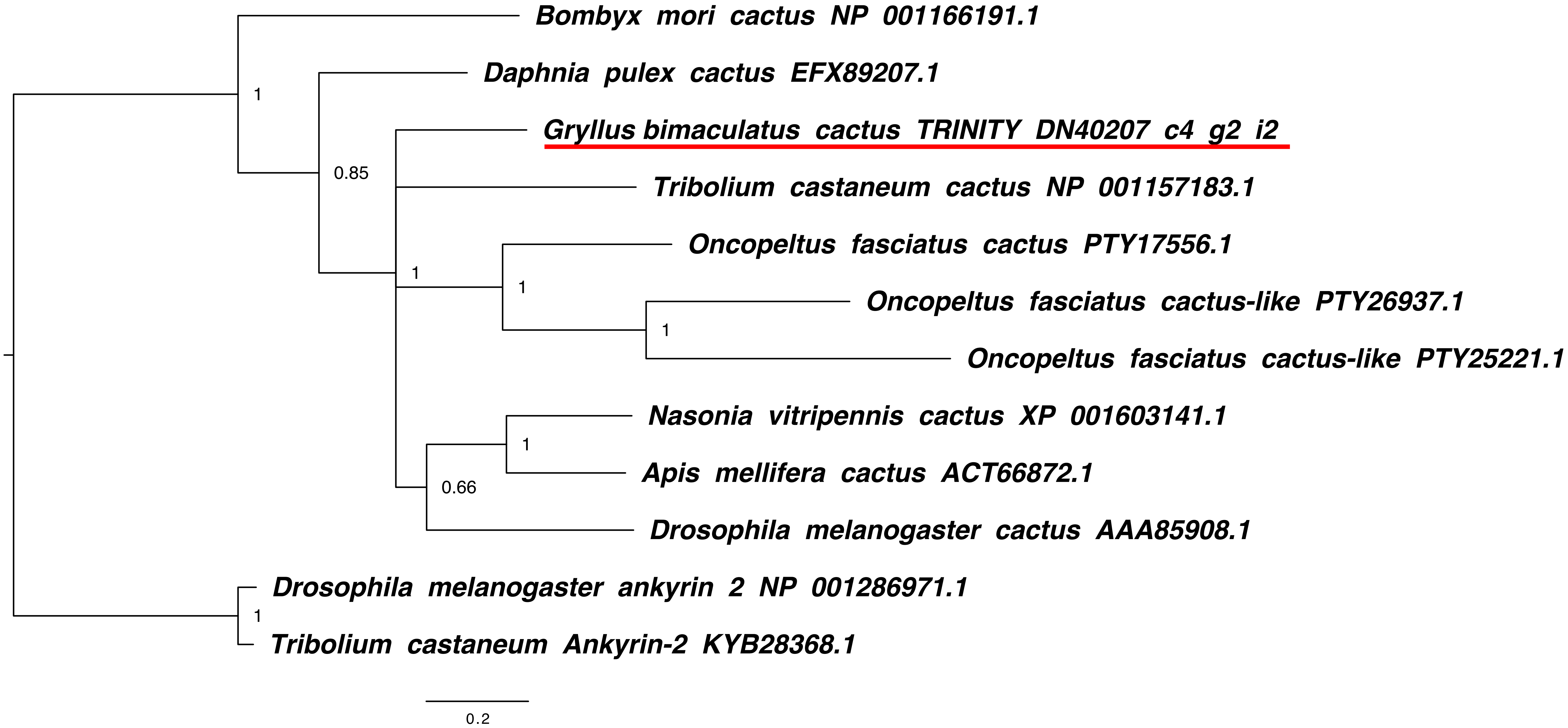

### cactus.png

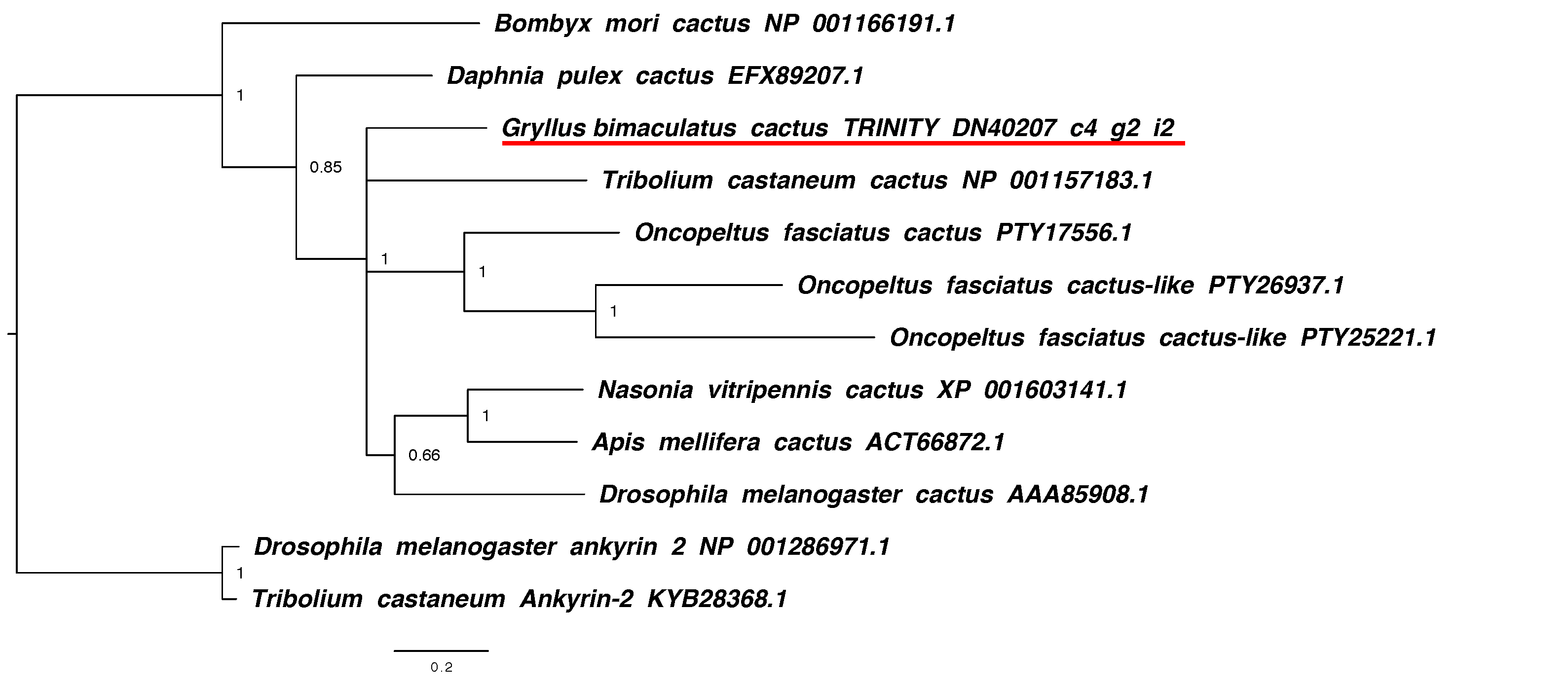

### Cerberus_Dan_Dante.pdf

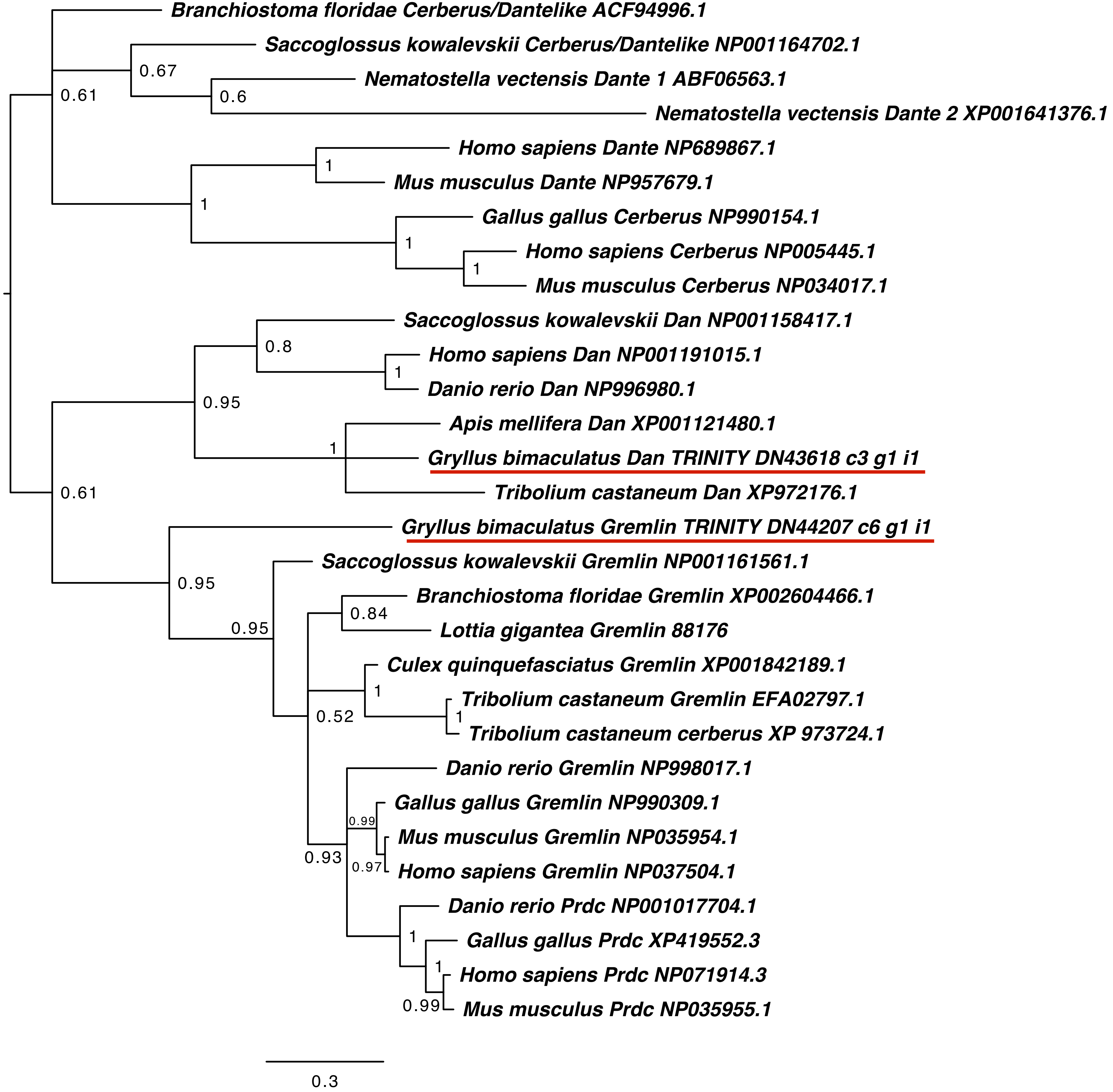

### dorsal.pdf

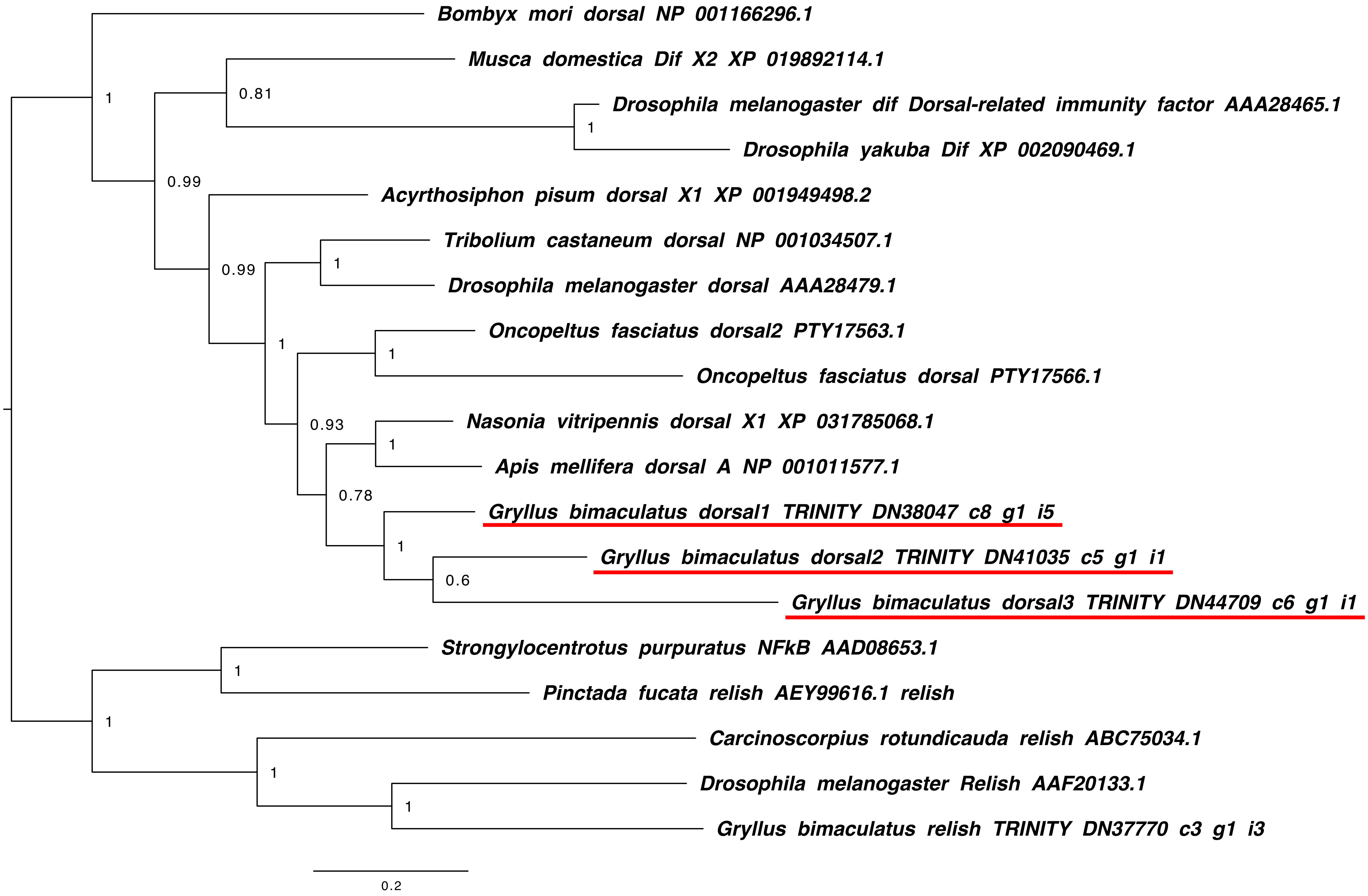

### follistatin.pdf

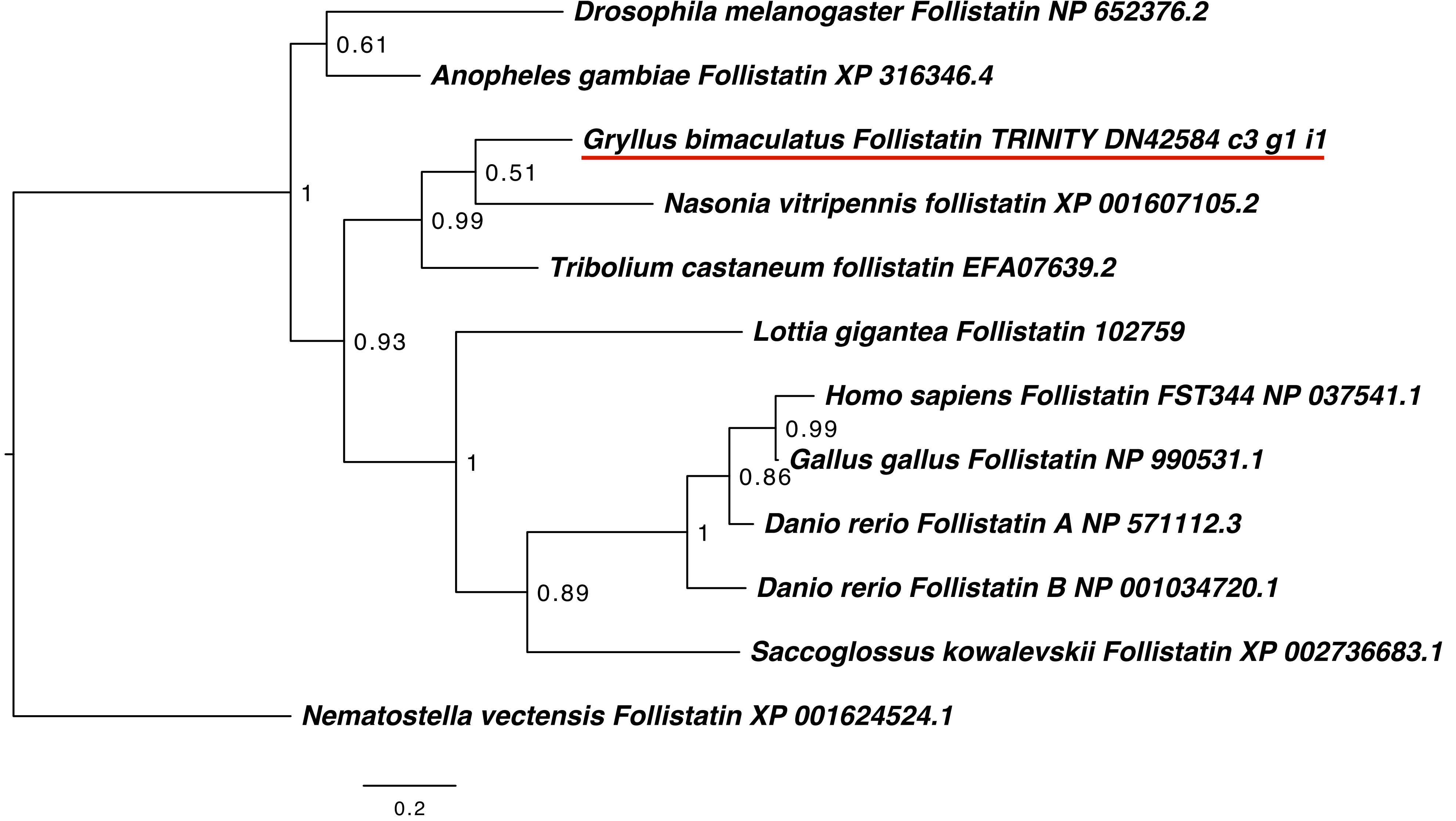

### KCP_BMPER_trim.pdf

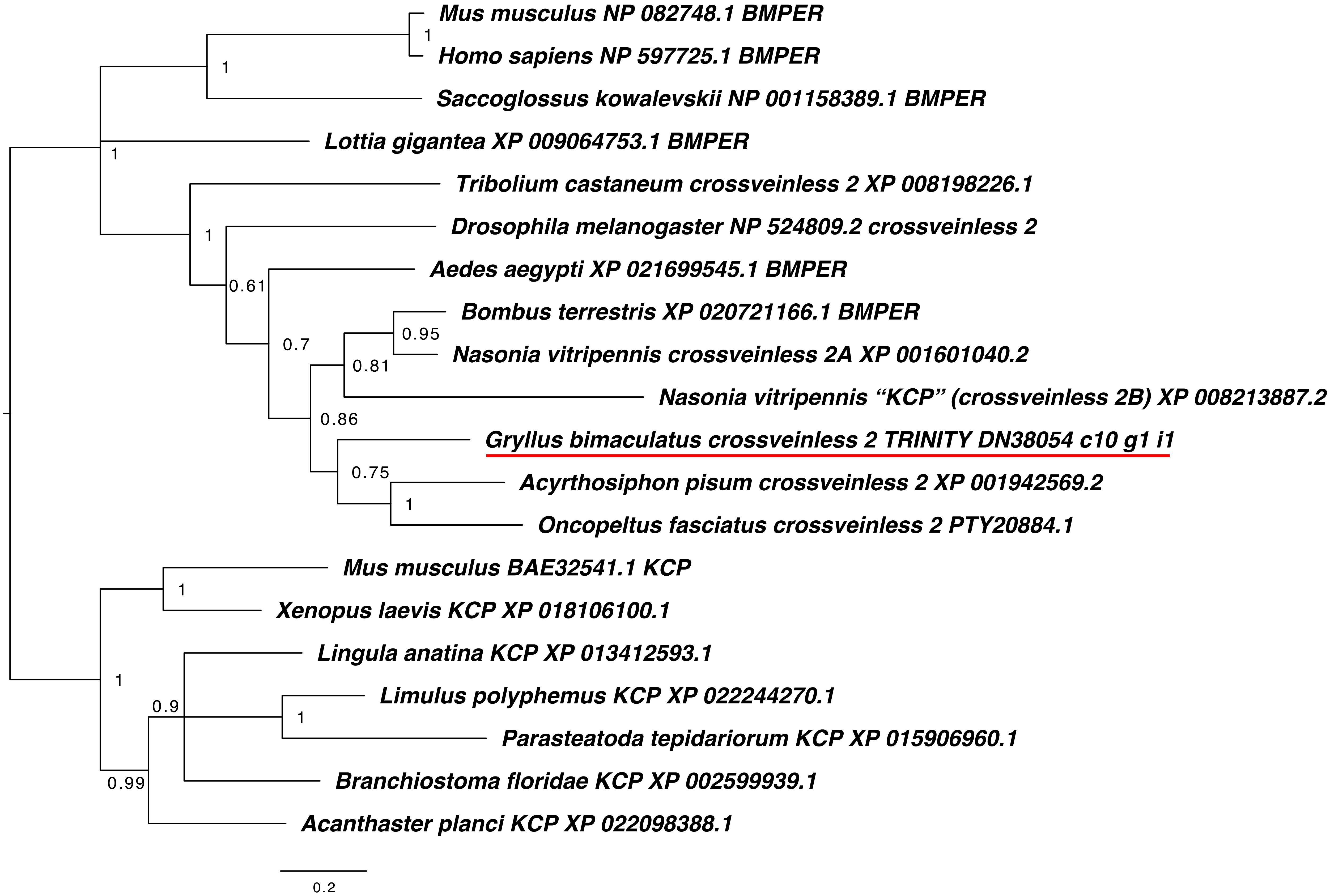

### ligands.pdf

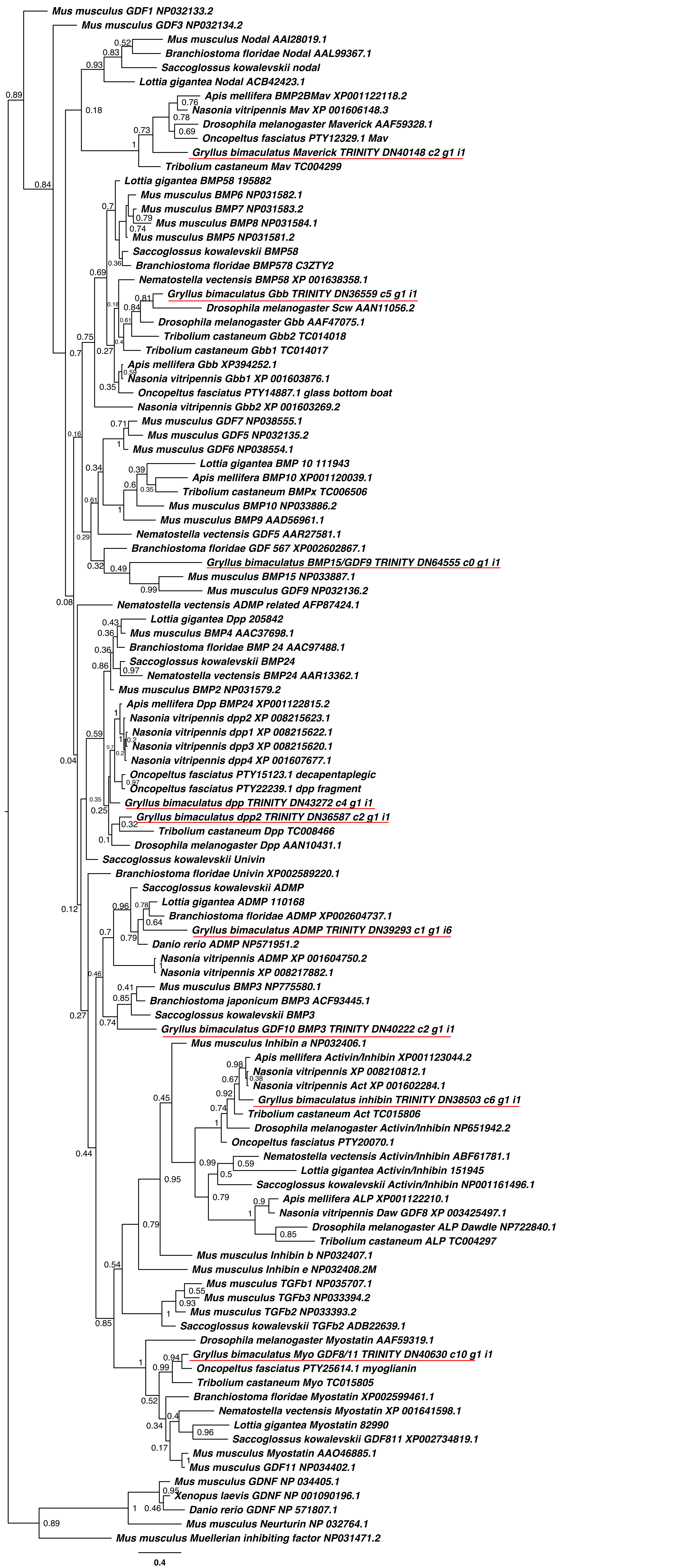

### MyD88.pdf

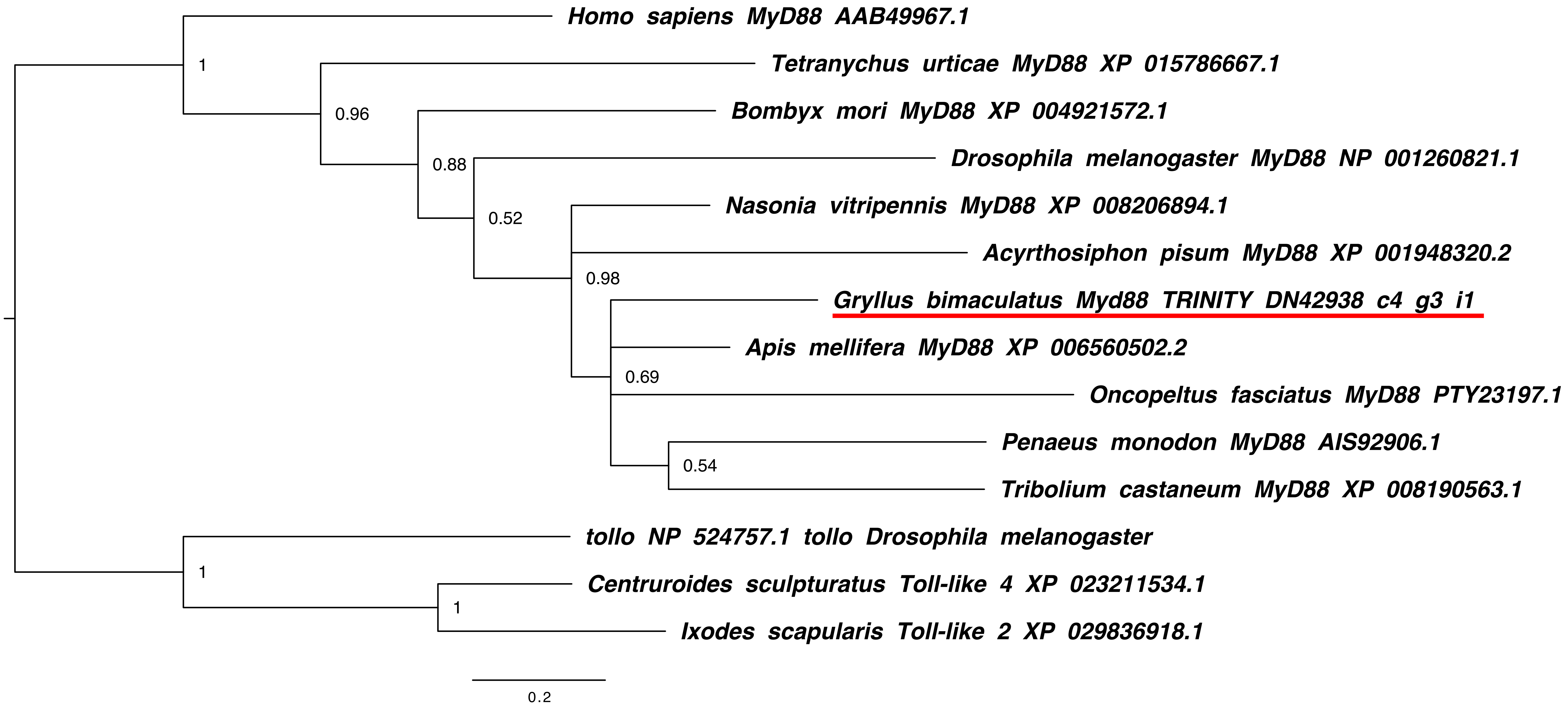

### MyD88.png

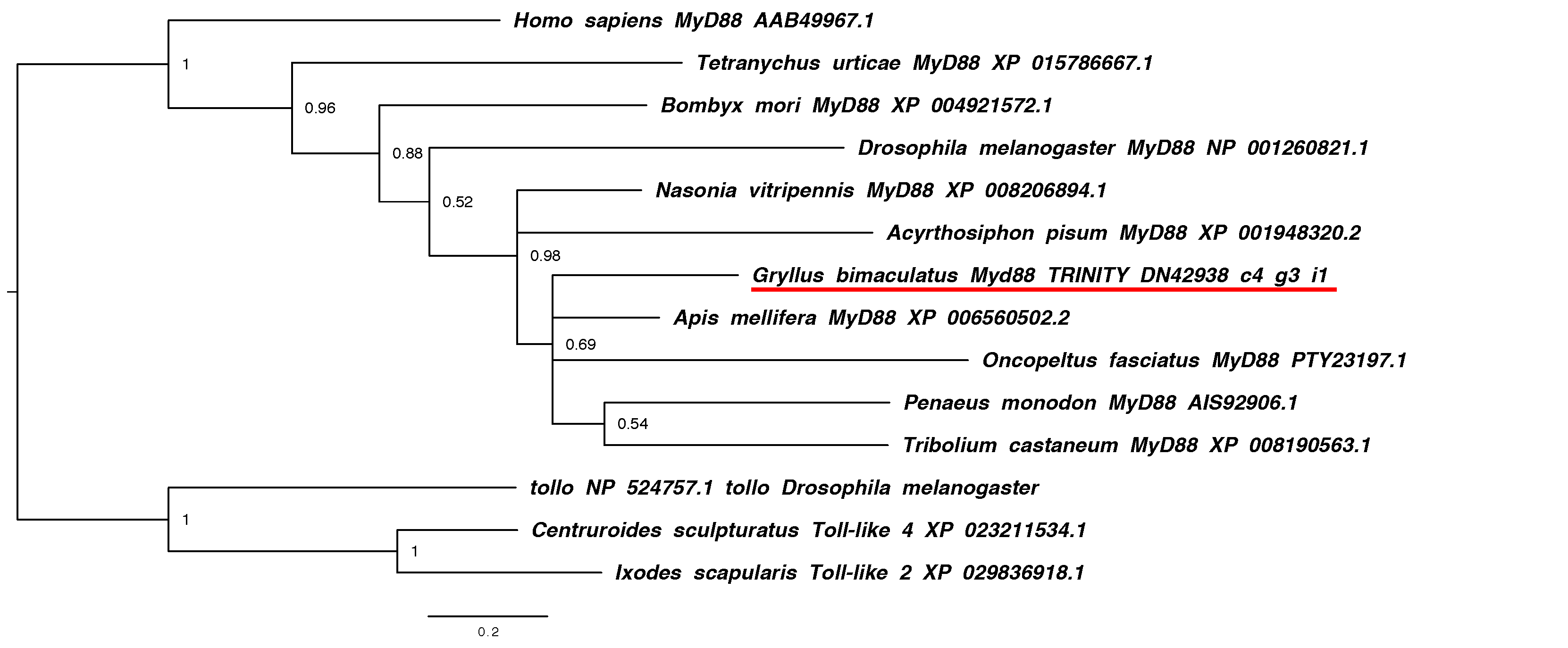

### noggin.pdf

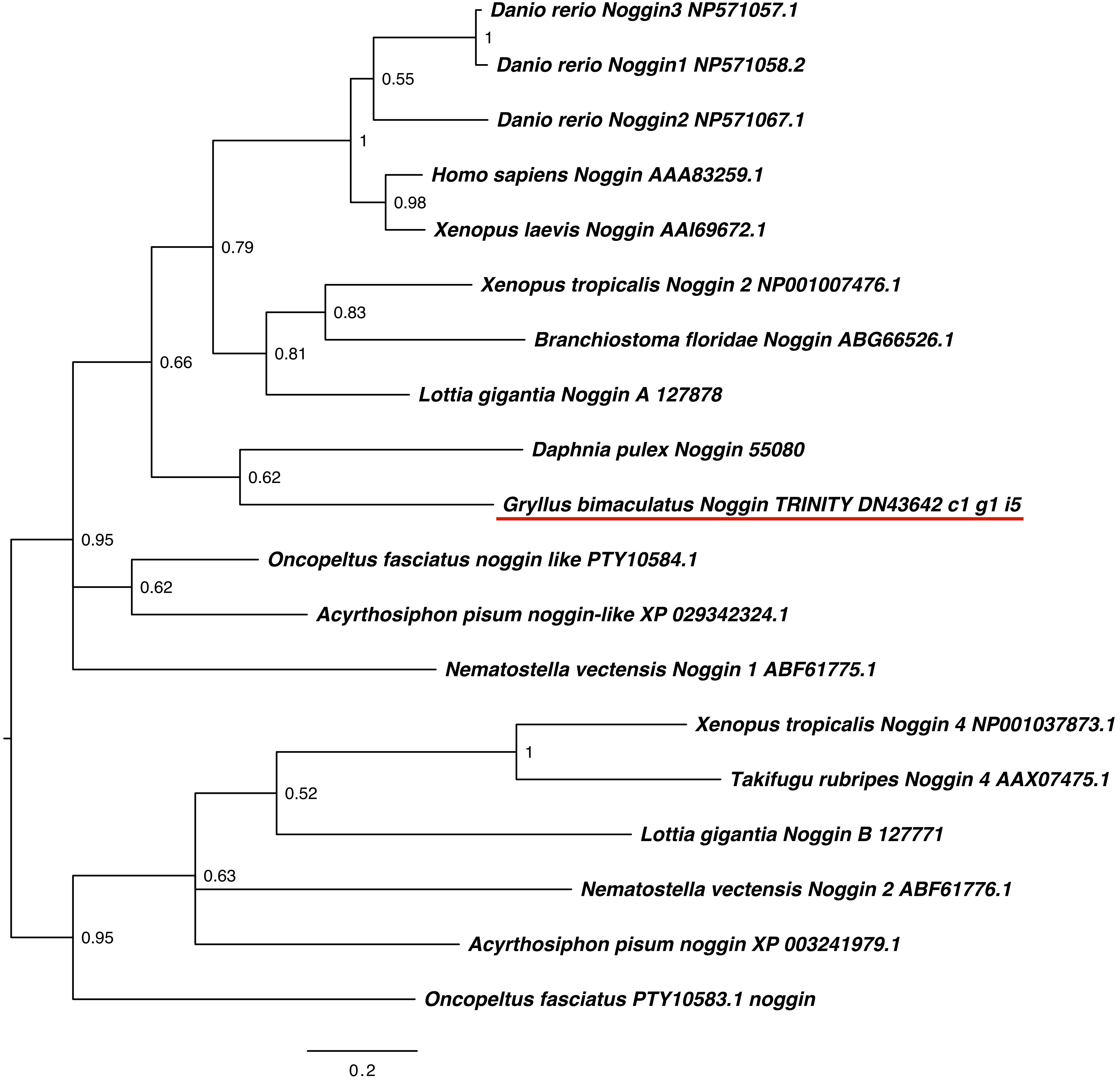

### nudel.pdf

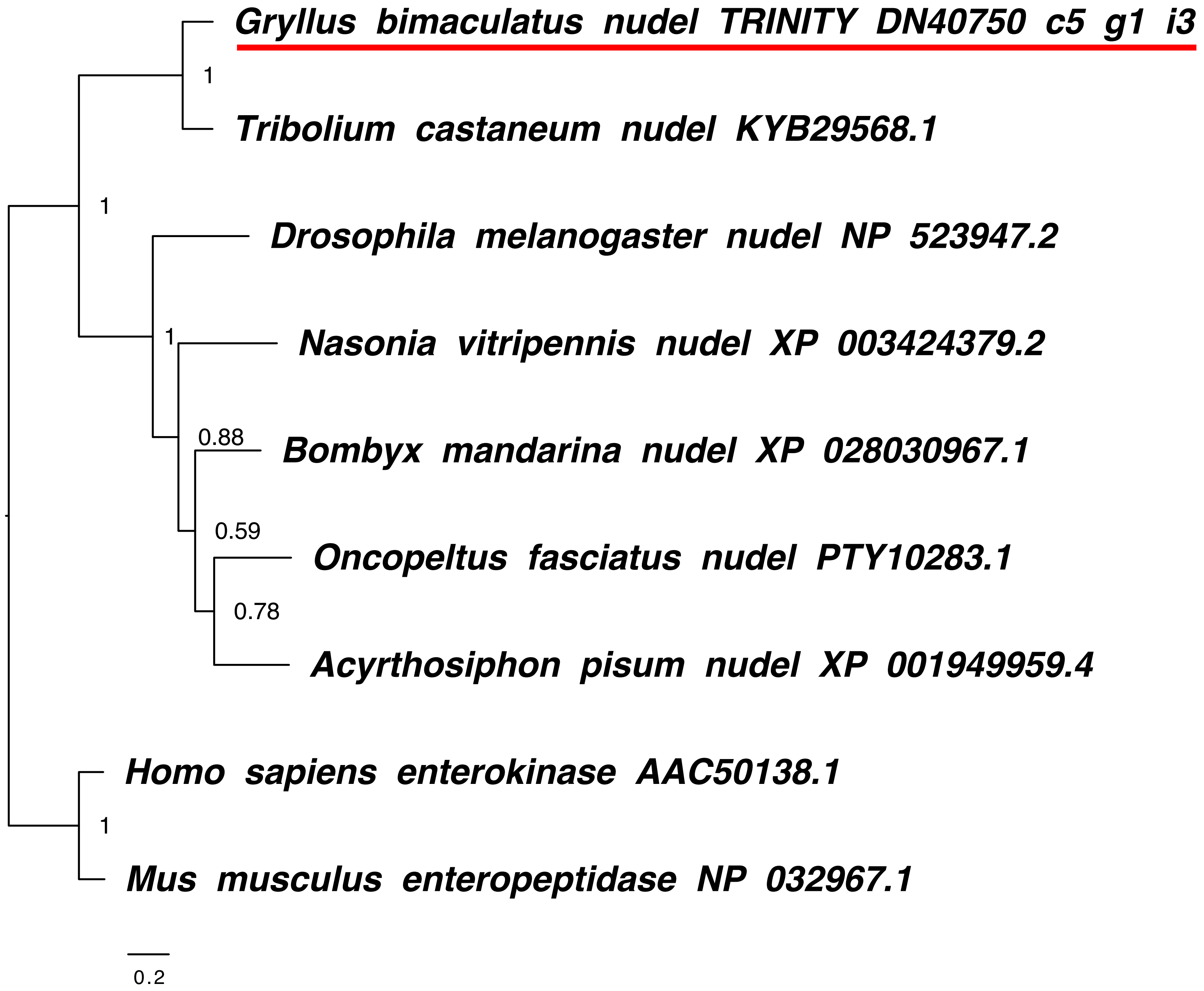

### nudel.png

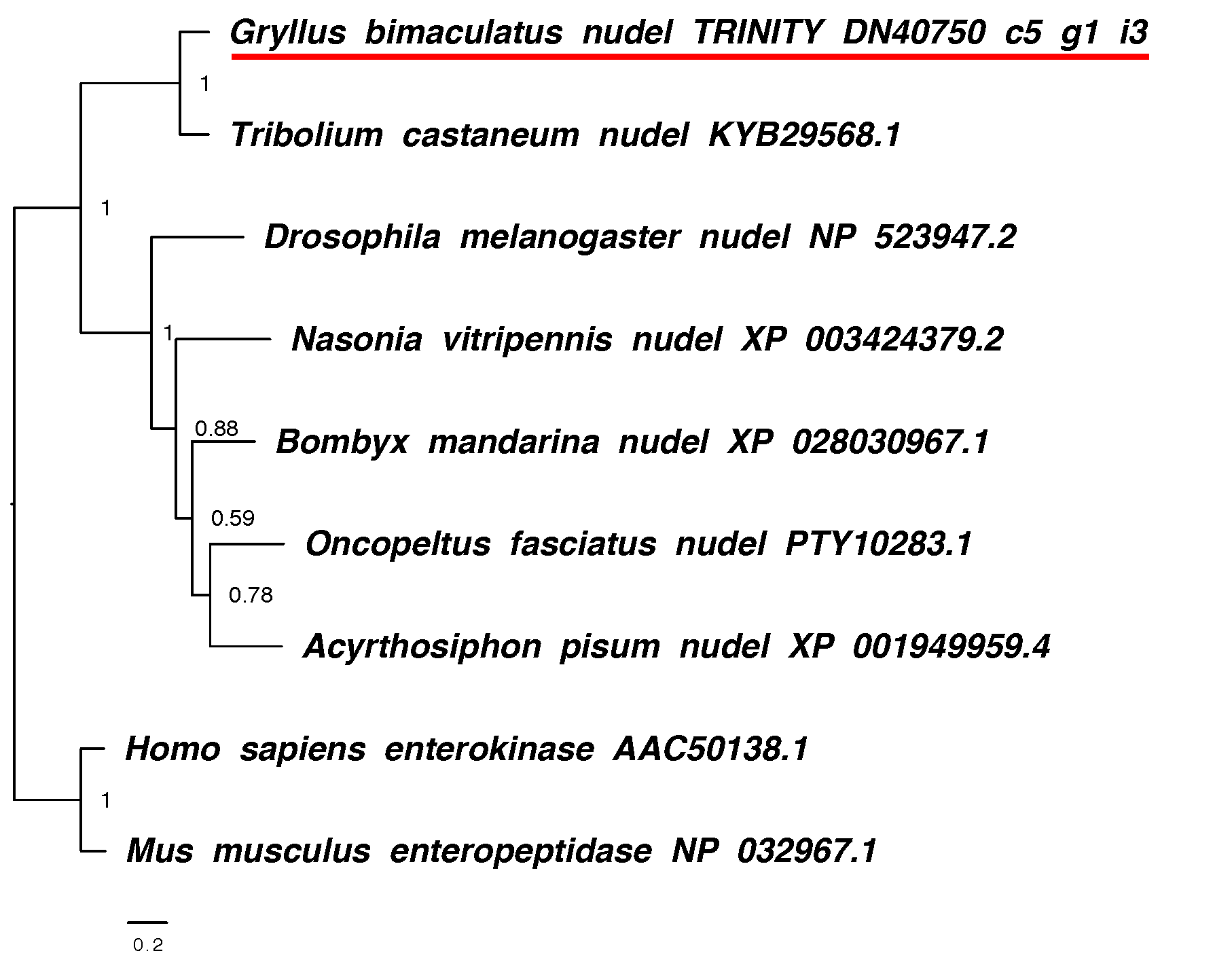

### pelle_tube.pdf

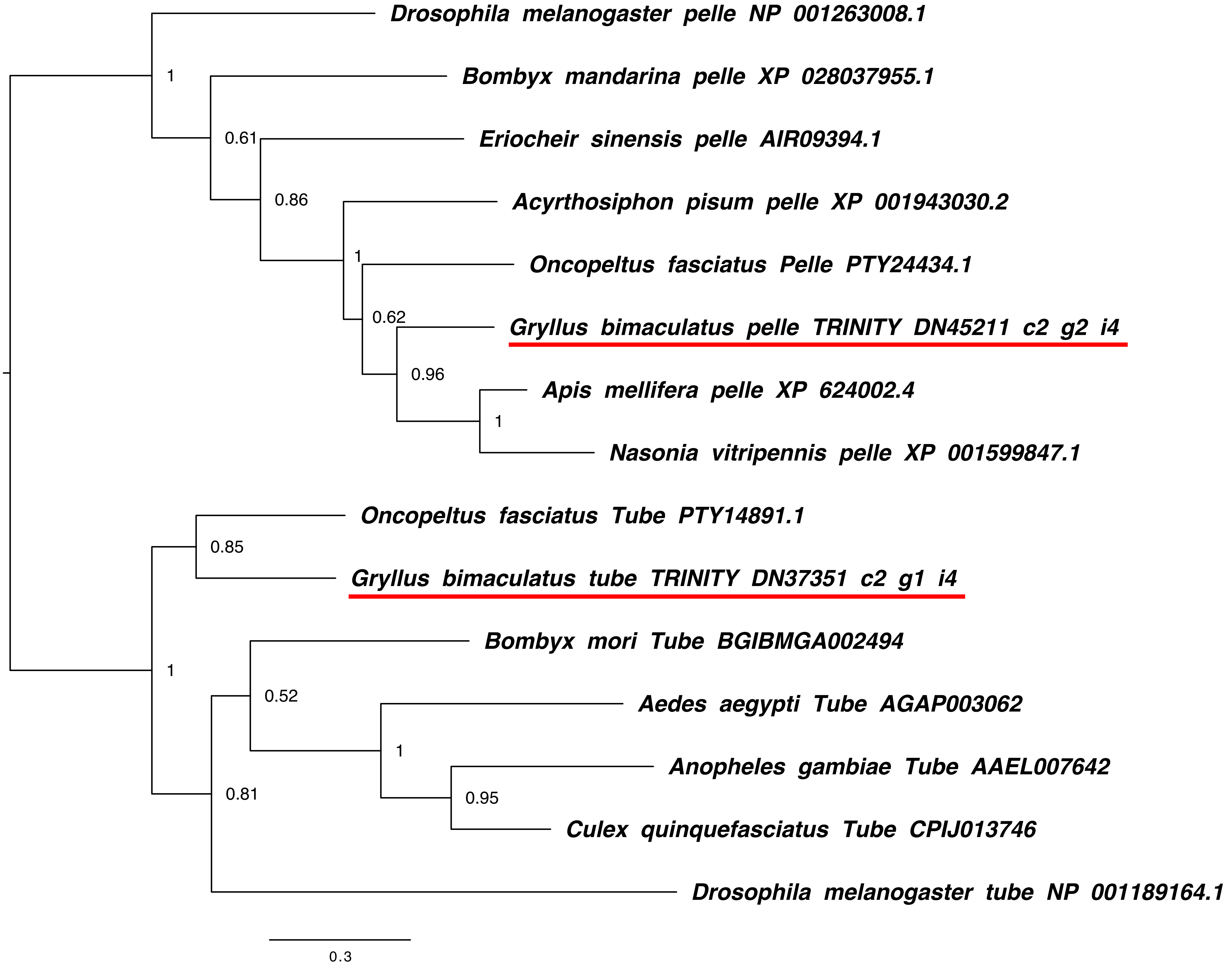

### pelle_tube.png

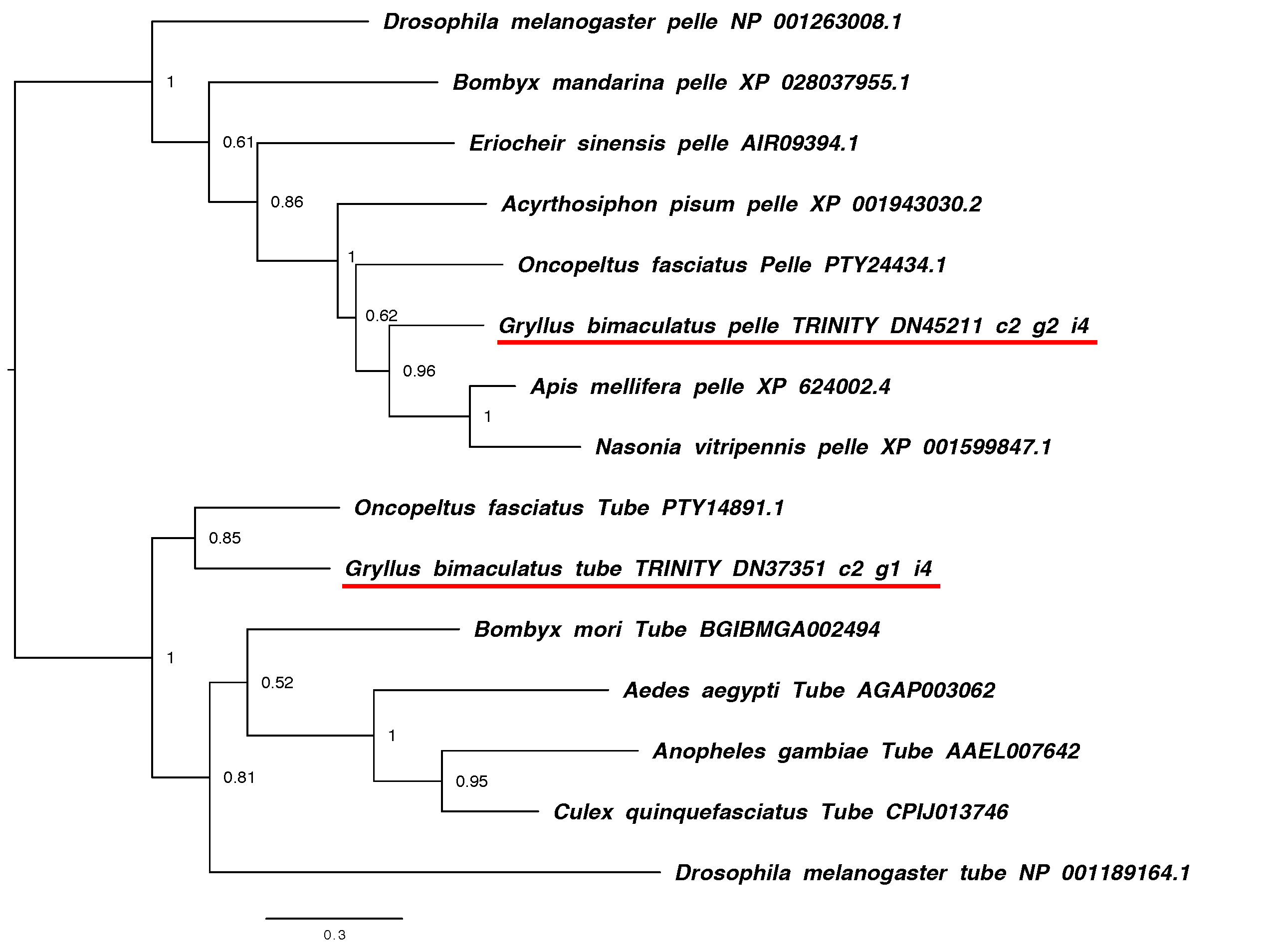

### pentagone.pdf

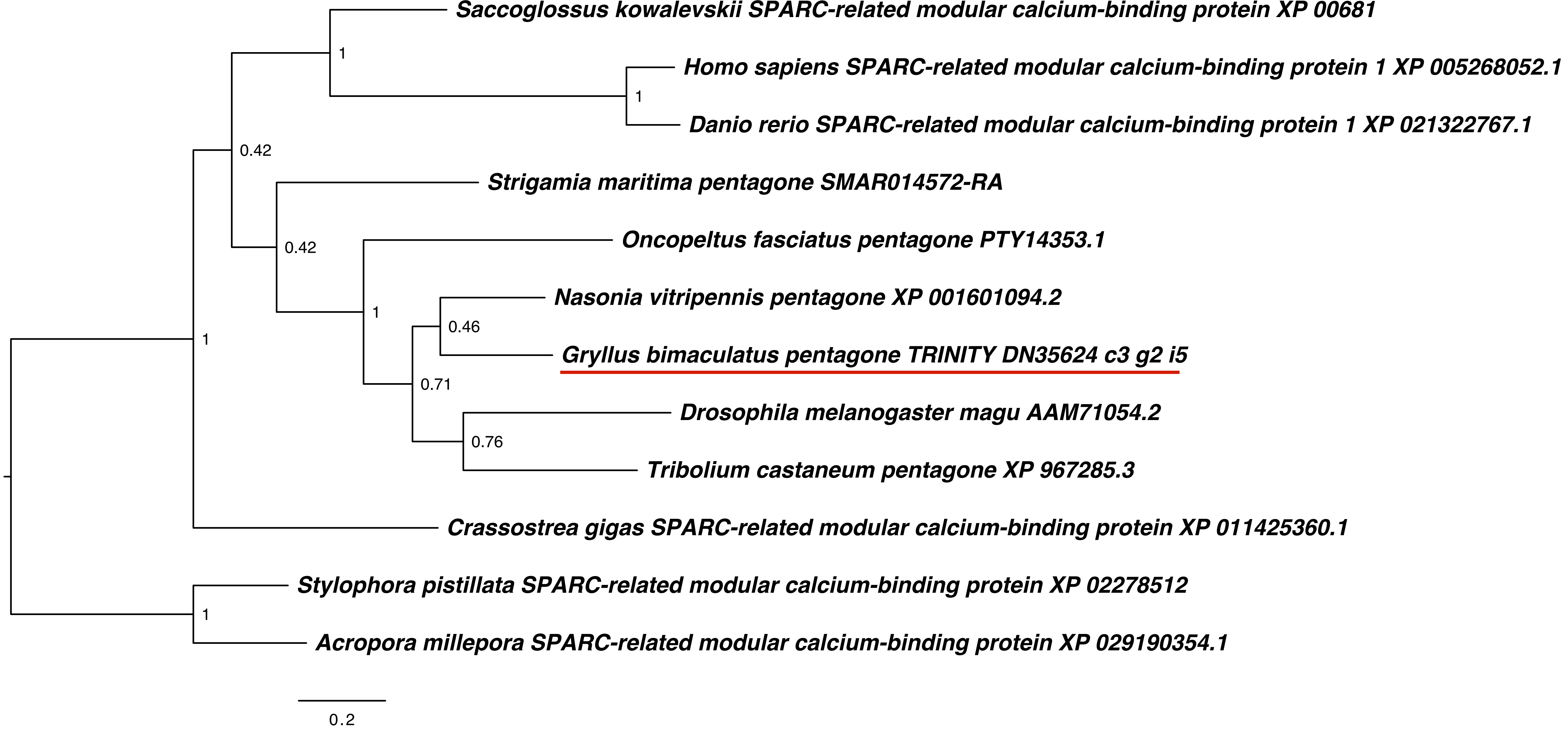

### pipe.pdf

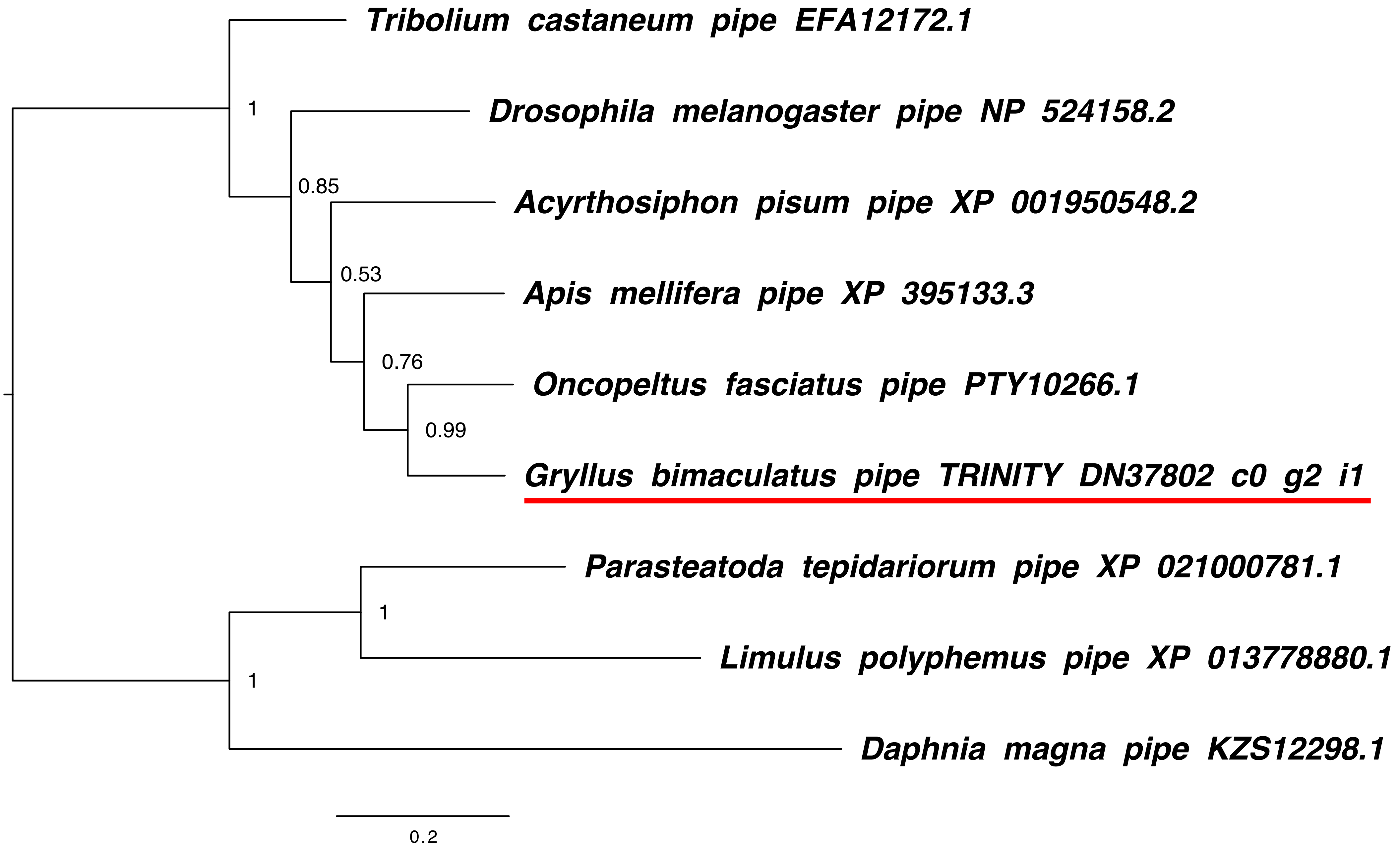

### pipe.png

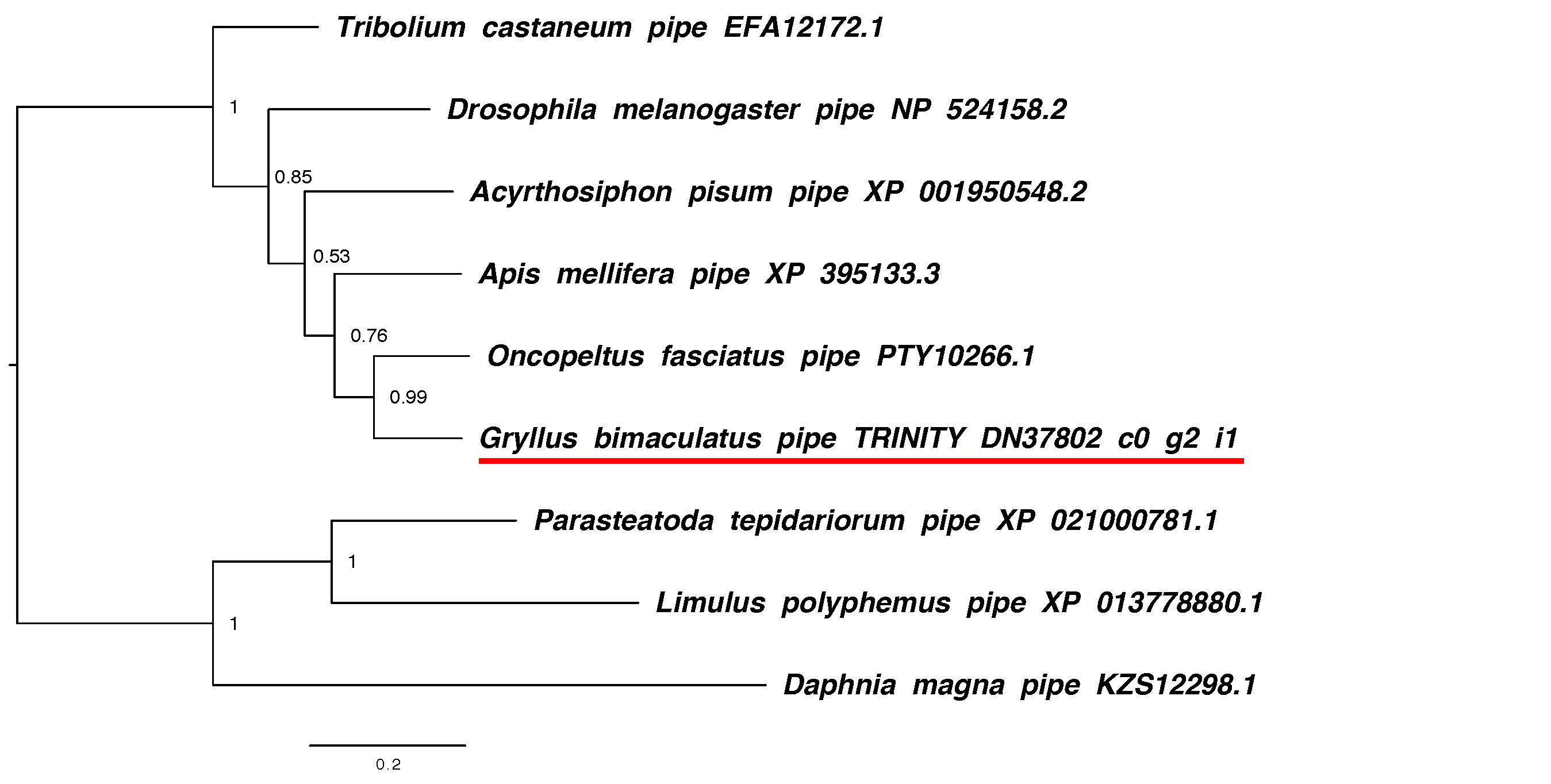

### schnurri.pdf

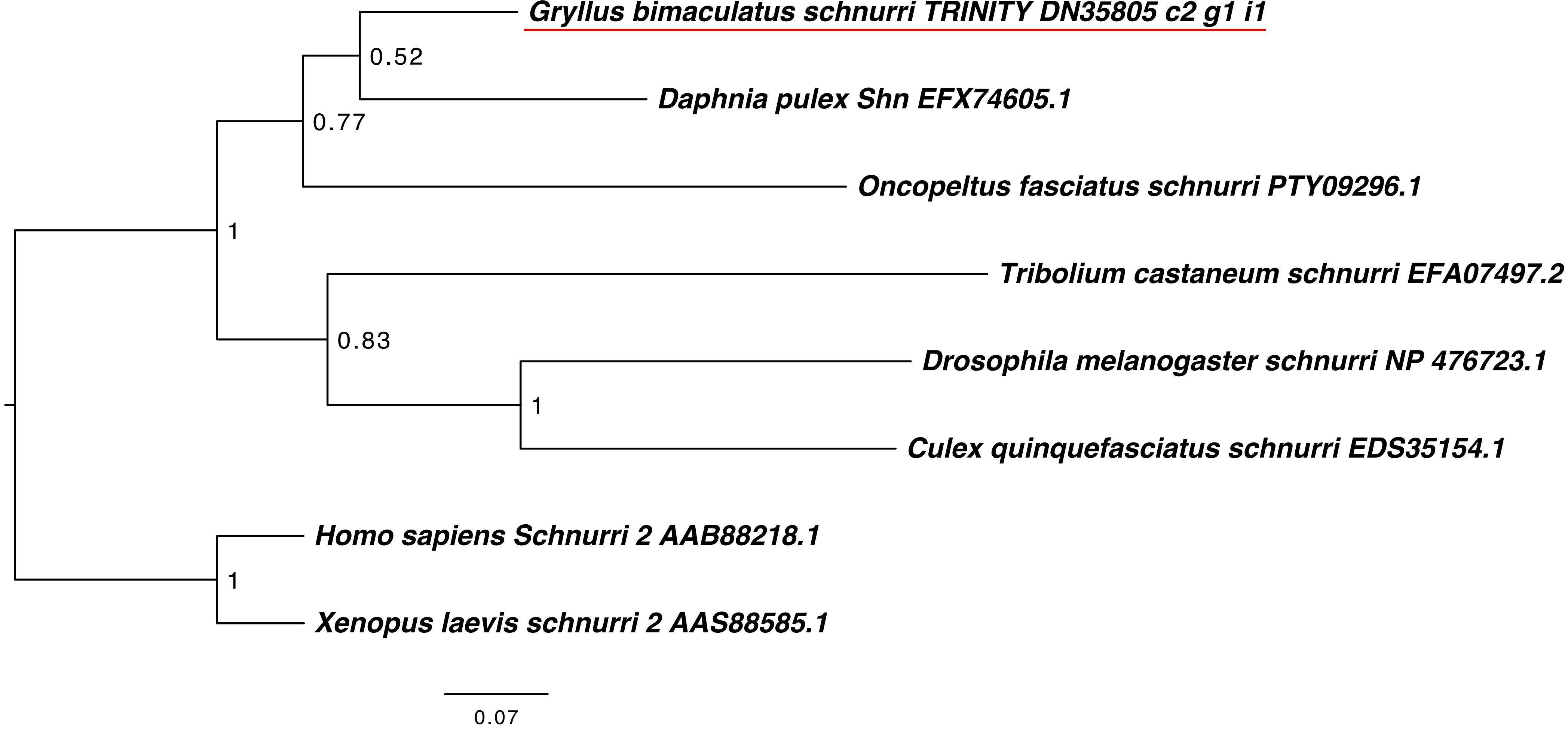

### smads.pdf

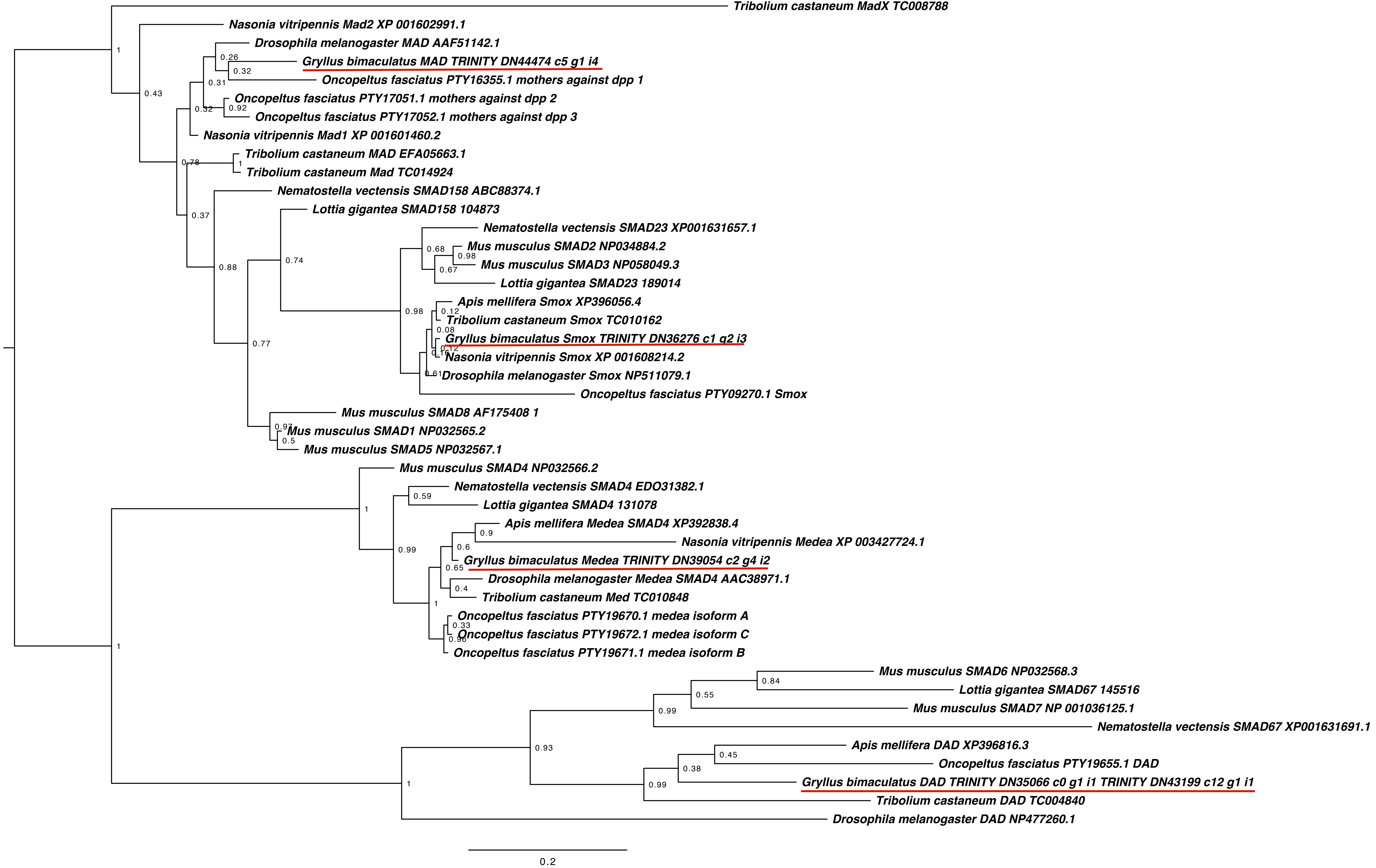

### smurf.pdf

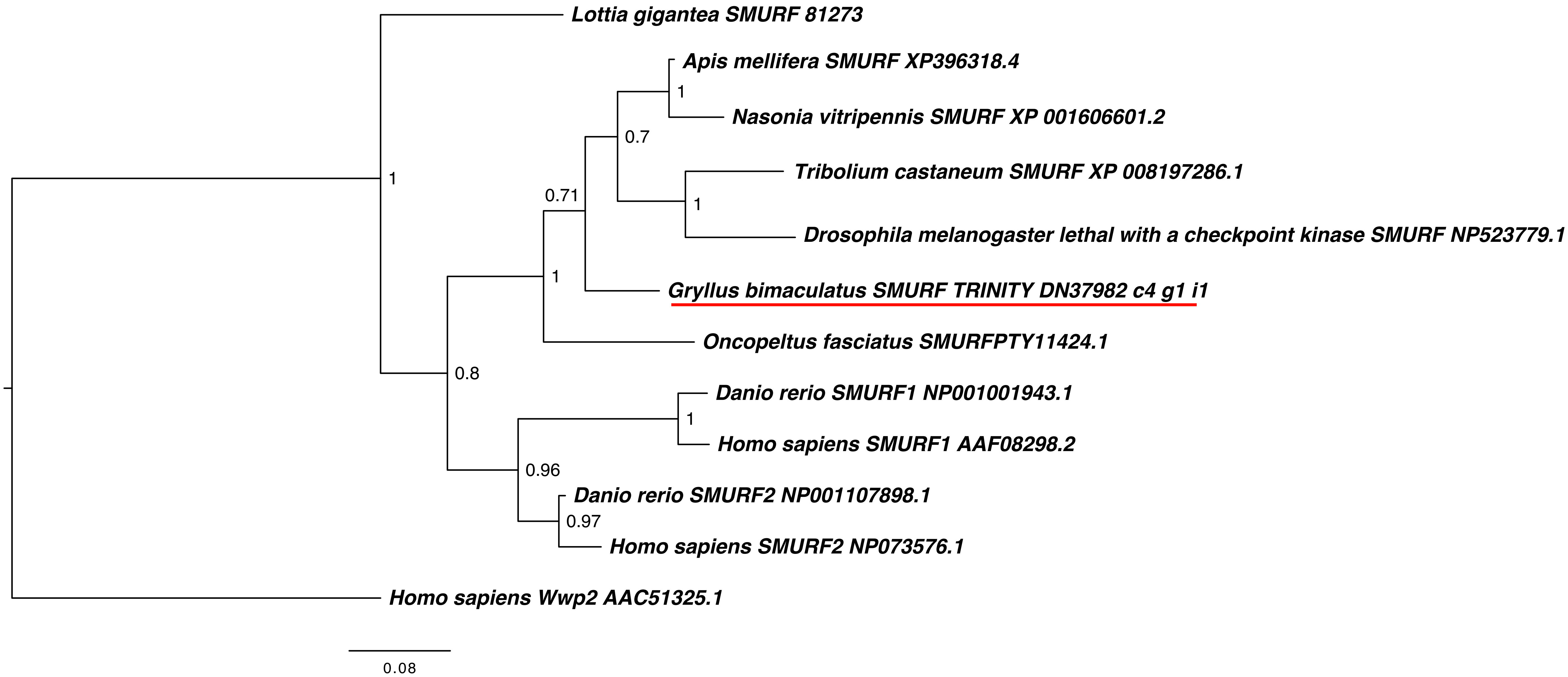

### spaetzle.pdf

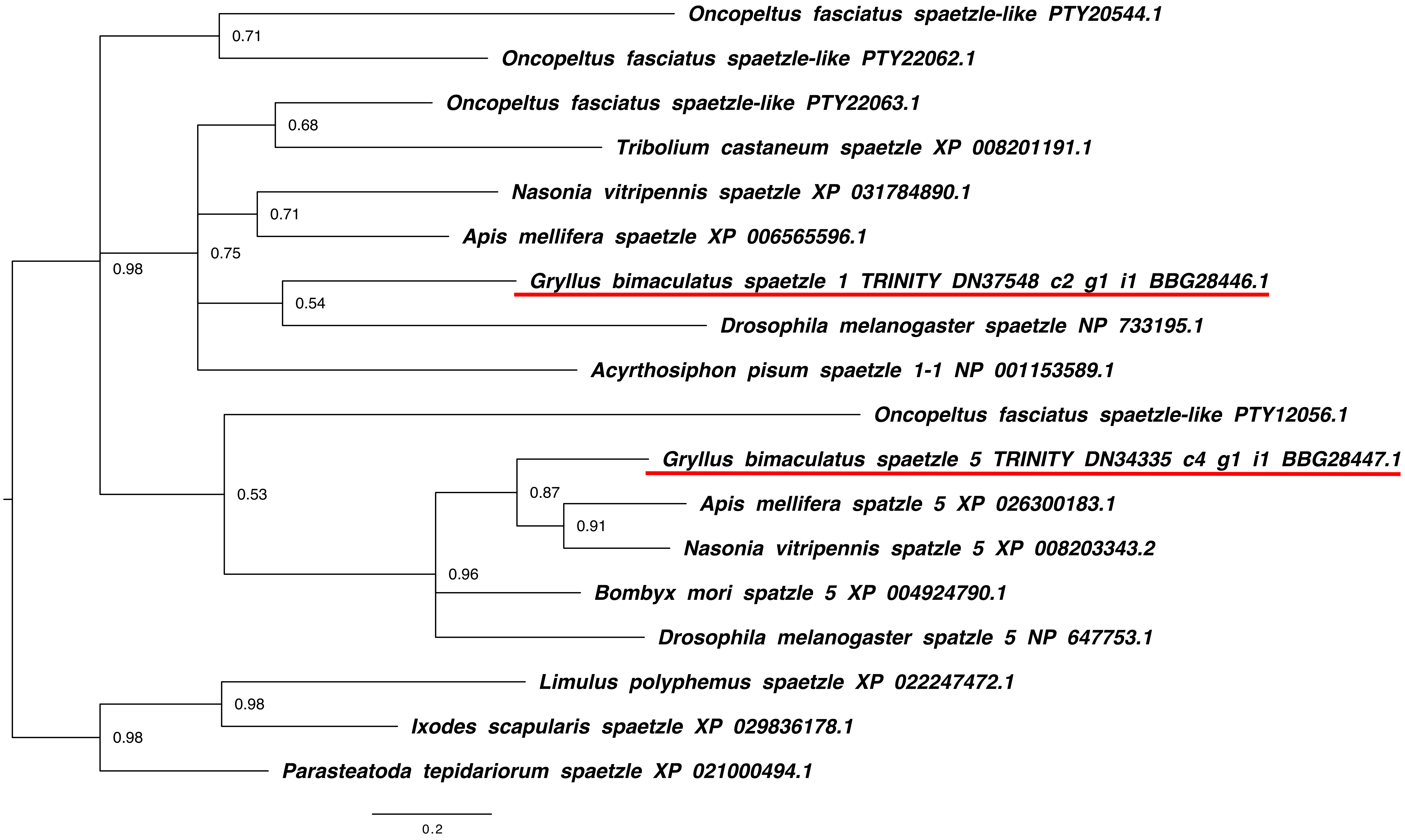
